# Single-nucleus sequencing reveals enriched expression of genetic risk factors in Extratelencephalic Neurons sensitive to degeneration in ALS

**DOI:** 10.1101/2021.07.12.452054

**Authors:** Francesco Limone, Daniel A. Mordes, Alexander Couto, Brian J. Joseph, Jana M. Mitchell, Martine Therrien, Sulagna Dia Ghosh, Daniel Meyer, Yingying Zhang, Melissa Goldman, Laura Bortolin, Inma Cobos, Irena Kadiu, Steven A. McCarroll, Beth Stevens, Olli Pietiläinen, Aaron Burberry, Kevin Eggan

**Affiliations:** Department of Stem Cell and Regenerative Biology, Harvard University, Cambridge, MA, USA; Stanley Centre for Psychiatric Research, Broad Institute of MIT and Harvard, Cambridge, MA, USA; Leiden University Medical Center, LUMC, 2333 ZA Leiden, The Netherlands; Department of Pathology, Massachusetts general Hospital, Boston, MA, USA; Neuroscience Center, Helsinki Institute of Life Science, University of Helsinki, Helsinki, Finland; Boston Children’s Hospital, F.M. Kirby Neurobiology Center, Boston, MA, USA; Department of Genetics, Harvard Medical School, Boston MA, USA; Neuroscience Therapeutic Area, New Medicines, UCB Pharma, Braine-l’Alleud, Belgium; Department of Pathology, School of Medicine, Case Western Reserve University, Cleveland, OH, USA

## Abstract

Amyotrophic Lateral Sclerosis (ALS) is a fatal neurodegenerative disorder characterised by a progressive loss of motor function. The eponymous spinal sclerosis observed at autopsy is the result of the degeneration of extratelencephalic neurons, Betz cells (ETNs, Cortico-Spinal Motor Neuron). It remains unclear why this neuronal subtype is selectively affected. To understand the unique molecular properties that sensitise these cells to ALS, we performed RNA sequencing of 79,169 single nuclei from cortices of patients and controls. In unaffected individuals, we found that expression of ALS risk genes was significantly enriched in *THY1*^+^-ETNs and not in other cell types. In patients, these genetic risk factors, as well as genes involved in protein homeostasis and stress responses, were significantly induced in a wide collection of ETNs, but not in neurons with more superficial identities. Examination of oligodendroglial and microglial nuclei revealed patient-specific changes that were at least in part a response to alterations in neurons: downregulation of myelinating genes in oligodendrocytes and upregulation of a reactive state connected to endo-lysosomal pathways in microglia. Our findings suggest that the selective vulnerability of extratelencephalic neurons is partly connected to their intrinsic molecular properties sensitising them to genetics and mechanisms of degeneration.

**Graphical abstract and working model:** Our study highlights cell type specific changes in premotor/motor cortex of sporadic ALS patients. Specifically, we identify upregulation of synaptic molecules in excitatory neurons of upper cortical layers, interestingly correlating to hyperexcitability phenotypes seen in patients. Moreover, excitatory neurons of the deeper layers of the cortex, that project to the spinal cord and are most affected by the disease, show higher levels of cellular stresses than other neuronal types. Correspondently, oligodendrocytes transition from a highly myelinating state to a more neuronally engaged state, probably to counteract stressed phenotypes seen in excitatory neurons. At the same time, microglia show a reactive state with specific upregulation of endo-lysosomal pathways.

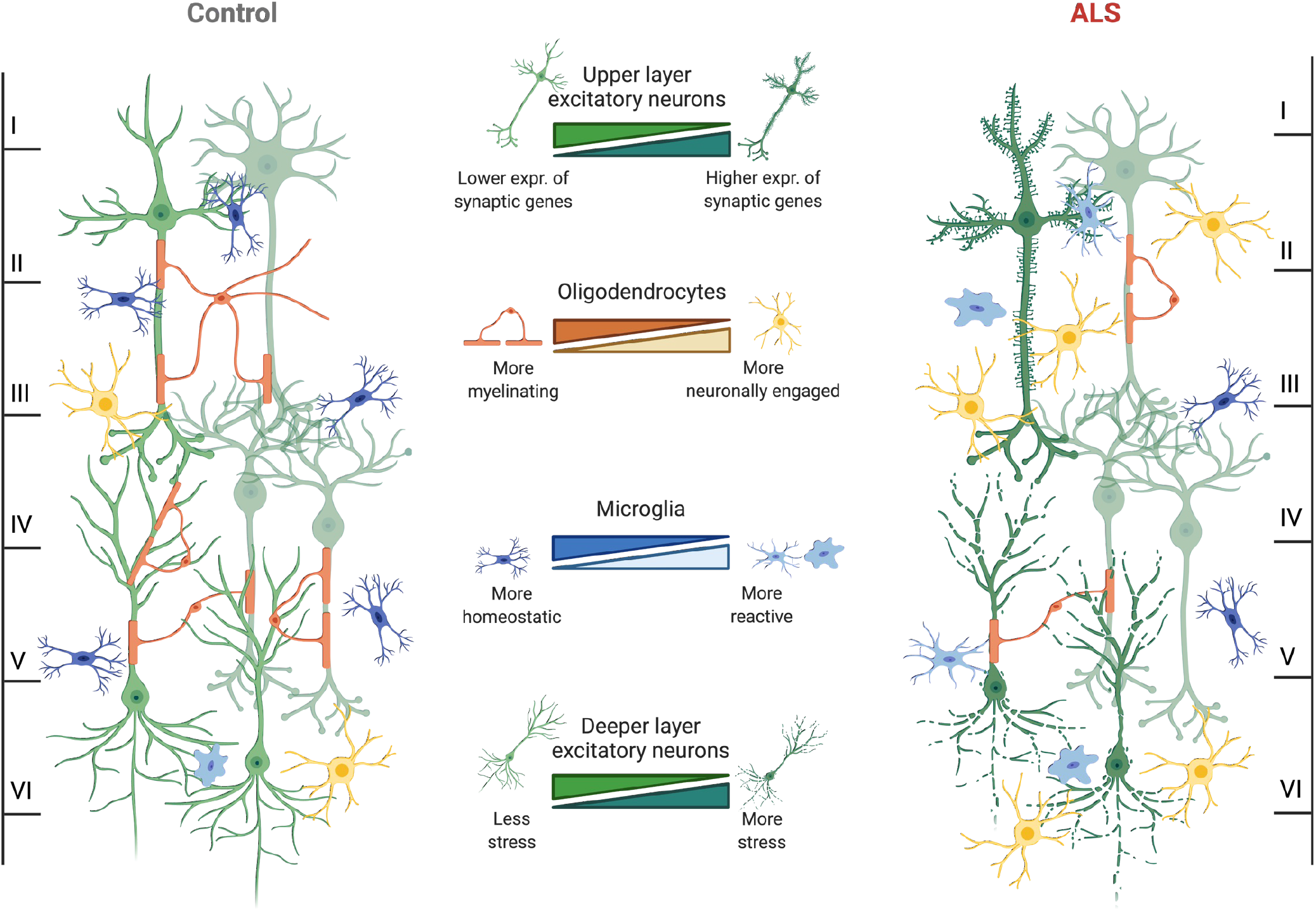

---

Amyotrophic Lateral Sclerosis (ALS) is a neuromuscular disease with survival typically limited to 2-5 years from onset, the most common motor neuron disease in aging individuals and the neurodegenerative disease with one of the earliest onsets, in the mid-to-late 50s^1^. Although specific genetic causes have been identified, most cases are sporadic (~90%), have no family history and unknown etiology^2^, rendering modelling of non-genetic forms of the disease difficult^3^. Variants in genes associated with ALS can contribute to a related disorder, Frontotemporal Dementia (FTD), leading to the view of ALS and FTD as different clinical manifestations of shared molecular causes. Bulk RNA-sequencing of ALS post-mortem brains have identified differences^4,5^ and similarities between sporadic and familial^6^ cases and highlighted shared profiles independent of disease onset^7,8^. While they have provided valuable insights, these studies have had limited resolution on the cell types mostly altered by the disease.

The most striking feature in ALS/FTD is the formation of protein aggregates of TAR DNA/RNA-binding protein-43 (TDP-43) in over 95% of cases of ALS and ~50% of FTD cases, mostly in neurons^9^, providing at least one shared mechanism. While the pattern of degeneration is similar, it is still unknown how familial mutations and sporadic onset might converge on the formation of these aggregates and how it specifically affects classes of extratelencephalic Cortico-Spinal Motor Neurons, i.e. Betz^10,11^ and von Economo cells^12,13^. Moreover, strong evidence demonstrated that cells other than neurons are key mediators of disease progression and it remains unclear how these might contribute to the disease^14–17^.

Methods to study heterogeneity at a single-cell level have rapidly advanced and their application to human post-mortem brain tissue is beginning to emerge for neurodegenerative diseases^18–27^. However, a comprehensive view across cell types in ALS has not been performed. In this study, we applied single-nuclei RNA sequencing and *in vitro* human induced Pluripotent Stem Cells modelling to investigate specific changes in cortical cell types in sporadic ALS. Our profiling identified the intrinsically higher expression of ALS/FTD risk factors in a specific class of extratelencephalic excitatory neurons. In ALS patients, these neurons selectively express higher levels of genes connected to unfolded protein responses and RNA metabolism. We also found that excitatory neurons vulnerability is accompanied by a decrease in myelination-related transcripts in oligodendroglial cells and an upregulation of reactive, pro-inflammatory state in microglial. We provide a preliminary, insightful view of disruptions triggered in human motor cortices in ALS and implicate aging-associated mechanisms like altered proteostasis, inflammation and senescence to specific cell type in the disease.

## Results

### Profiling of ALS cortex by single-nucleus RNA-sequencing

To better understand factors that contribute to the specific degeneration of classes of excitatory neurons, we used snRNAseq to profile motor/pre-motor cortex grey matter from sporadic (sALS) patients and age-matched controls with no neurological disease using Drop-seq technology^28^ (Fig. 1a, Extended Data Table 1, Extended Data Fig. 1a-c). After screening for RNA quality, 79,169 barcoded droplets from 8 individuals were analysed (*n*=5 sALS, *n*=3 Control), with a mean of 1269 genes and 2026 unique molecular identifiers (UMIs) (Extended Data Fig. 1d). We used Seurat^29^, single-cell analysis package, to cluster and annotate groups according to canonical markers of brain cell types^30^: excitatory and inhibitory neurons, oligodendrocytes, oligodendrocyte progenitor cells (OPCs), microglia, astrocytes, and endothelial cells (Extended Data Figure 1e,f). The observed cell type distribution corresponded to previous studies^31^ and enabled robust categorization for downstream analysis. Cellular distribution was homogeneous between sexes and individuals, except for a modestly lower number of astrocytes in ALS samples (Extended Data Fig. 1g-i).

**Fig. 1.**
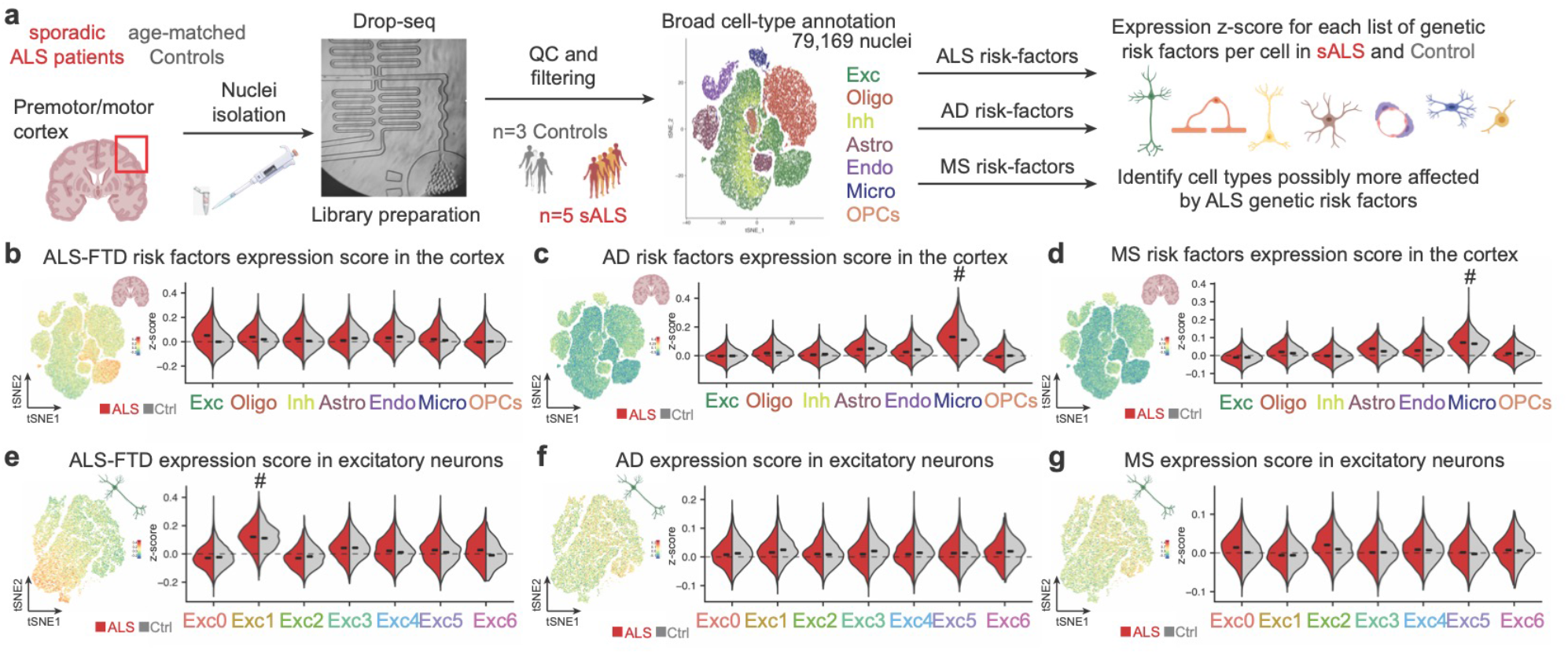
Cellular susceptibility to ALS-FTD in the human cortex. **a**, Diagram of workflow for isolation of nuclei from cortices of ALS patients and age-matched controls followed by single-nucleus RNA sequencing and assessment of expression of gene modules associated with neurodegenerative diseases. **b-d**, *t*-SNE projections and Violin plots of z-scores for expression of genes associated with the ALS-FTD (b), AD (c) and MS (d) in the different cell types identified (bars denote median for each side of the violin plot). (#) statistically significant differences. **e-g**, *t*-SNE projections and Violin plots of z-scores for expression of genes associated with the ALS-FTD (e), AD (f) and MS (g) in the different subtypes of excitatory neurons (bars denote median for each side of the violin plot). (#) statistically significant differences.

### Elevated expression of ALS/FTD risk genes in a specific class of excitatory neurons

To potentially identify cell types underlying ALS pathophysiology, we examined the expression of known familial genes for ALS/FTD and variants identified as risk factors from genome-wide association studies (GWAS). These genes were expressed to a highly variable degree between cell types and many of them were ubiquitously expressed as already known in the field^2^ (Extended Data Fig. 2a). We then computed a “module score” for this set of genes^32^; this metric generates a standardised z-score for the expression of each gene and sums it up as a total score for the gene set, here a positive score suggests higher expression of this gene set compared to the average expression of the module across the dataset. We also computed parallel module scores for lists compiled from latest GWAS for neurological disorders that affect the cortex but not specifically Betz cells: AD^33,34^ and MS^35^ (Fig. 1a, Extended Data Table 2). No clear preferential expression for ALS/FTD gene list was identified (Fig. 1b), as it might have been anticipated by the scattered and ubiquitous pattern of expression. On the other hand, AD and MS modules showed enrichment for their respective lists in microglia, as expected based on the strong immune signature that characterises these diseases and the involvement of immune cells in neurodegeneration^33–37^, as shown by other reports^30^ (Fig. 1c,d). These results corroborate knowledge in the field, underlying the strength of this analysis, and confirm our results in an unbiased, single-cell resolution.

Considering the selective loss of neurons in ALS, we further analysed these cells. We found 32,810 nuclei from excitatory neurons with unbiased clustering identifying seven subgroups (Exc0-6) that expressed known markers of different cortical layers, equally distributed in our cohort (Extended Data Fig. 2b-f). Analysis of the ALS/FTD module in these cells showed a positive score in *THY1*-expressing subgroup Exc1 (Normalised Enrichment Score=1.834) (Fig. 1e, Extended Data Fig. 2g,h) and no significant enrichment for AD and MS modules (Fig. 1f,g). We decided to further dissect the identity of these cells and investigate if they could be ETNs.

We identified three subgroups expressing markers of subcerebral projection neurons: Exc1, Exc5 and Exc6 (Fig 2a). Exc5 and Exc6 expressed canonical markers *FEZF2, BCL11B* and *CRYM^38^*; Exc1 expressed *THY1*, enriched in human layer V^18^ and used as a reporter for CSMNs^39^, and high levels of neurofilament chains, markers of ETNs *in vivo^40^* (Fig. 2b). Recent reports dissected the transcriptomic identity of layer V extratelencephalic neurons in the human Motor Cortex^41^. We detected expression of their markers in these groups, with Exc1 expressing *SERPINE2* and *POU3F1*, specific of ETNs^41^, and *NEFH* and *STMN2*, broad markers of MN^40,42^ (Fig 2c). Because of the anatomical location of our samples and the presence of ETNs across motor-related areas^43^, we plotted markers specific to layer V ETNs of regions adjacent to the Motor Cortex like von Economo cells^44^, affected in FTD^45^, and other Long-Range Subcerebral Projecting Neurons (LR-SCPNs)^46^ and confirmed that all three groups expressed these markers (Fig. 2d,e). To further characterise the expression pattern of these markers we leveraged a publicly available single-cell, spatial dataset of the human dorsal cortex^47^. We confirmed that layer V markers expressed in Exc1 *THY1, STMN2* and *SNCG* (Fig. 2f) are specifically expressed in layer V (L5) in spatial dataset (Fig. 2g,h and Extended Data Fig. 3a,b). This evidence suggests that Exc1, Exc5 and Exc6 express markers of layer V extratelencephalic neurons of cortical areas affected by ALS/FTD.

**Fig. 2.**
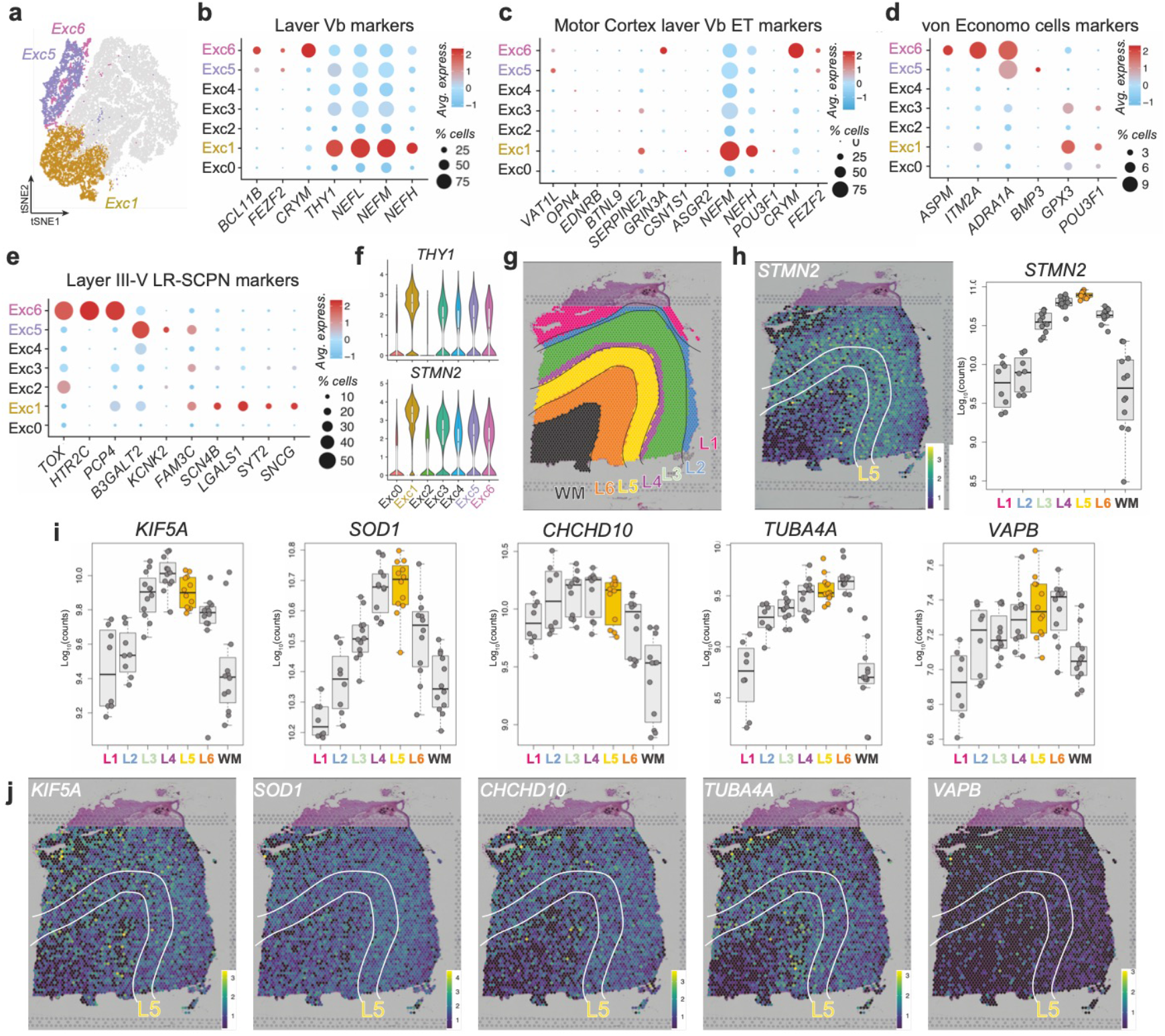
ALS-FTD susceptible neurons are layer V ET Neurons. **a**, *t*-SNE projection of presumptive layer V neurons. **b**, Dotplot representing expression of Layer V markers **c**, Dotplot for markers of LVb Extratelencephalic neurons of human Motor Cortex. **d**, Dotplot representing expression of von Economo markers. **e**, Dotplot representing expression LR-SCPN markers. **f**, Representative Violin for markers of layer V Extratelencephalic neurons of human Motor Cortex. **g**, Visual depiction of layers identification by Maynard et al. 2021 (publically available). **h**, Spotplot depicting expression of layer Vb Motor Cortex marker, *STMN2*, identified as enriched in THY1-Exc1, with corresponding boxplot quantification. **i-j**, Spotplot and corresponding boxplot for the expression of top 5 ALS/FTD associated genes expressed in Exc1.

To further confirm that *THY1*^high^-neurons expressed higher levels of ALS/FTD genes, we ran module score analysis in two datasets that identified *THY1*^high^ cortical neurons^18,48^. In these studies, *THY1*-neurons expressed ETNs markers, layer V, von Economo and LR-SCPNs markers (Extended Data Fig. 3c-i) and, expressed higher levels of the ALS/FTD module score (Extended Data Fig. 3l-m). Analysis of the spatial transcriptomic dataset^47^, confirmed that the top 10 ALS/FTD-associated genes most high expressed in Exc1 (Extended Data Fig. 2g) are highly expressed in deeper layers of the cortex and specifically in layer V (Fig. 2i,j and Extended Data Fig. 3n). Studies in human^13,49^ and mouse^50^ showed that deep layer neurons have a higher propensity to form TDP-43 aggregates, hallmark of ALS/FTD. Here we provide a possible link to their specific vulnerability.

### Cellular burden on excitatory neurons is higher in deeper layers

We next examined how the enriched expression of ALS/FTD genes relates to changes that occur in excitatory neurons in response to ALS. We conducted differential gene expression (DGE) analysis between neurons from patients and controls, across all excitatory cells and within each subgroup (Fig. 3a). We then selected genes significantly upregulated in patients globally (DGEall) and within each subgroup (DGE0-6), calculated module-scores for each set and investigated whether certain neuronal subtypes might have similar responses to ALS (Extended Data Table 3). This analysis showed a correlation between scores in groups expressing deep layer markers and the global changes identified in patients (Fig. 3b), suggesting that pathology in lower cortical layers are driving the observed alterations. For instance, groups expressing ETNs markers (Exc1, Exc5, Exc6) shared many upregulated genes with each other and with the global signature (Fig. 3c), whereas genes upregulated in upper layers of the cortex, a region relatively spared of pathology, shared less similarities (Fig. 3d). Intriguingly, this class of genes is, like genetic risk factors, constitutively expressed at higher levels in Exc1-ETNs (Fig. 3b), advocating for a proposed interplay between genetics and molecular pathways that sensitises ETNs to ALS^51^.

**Fig. 3.**
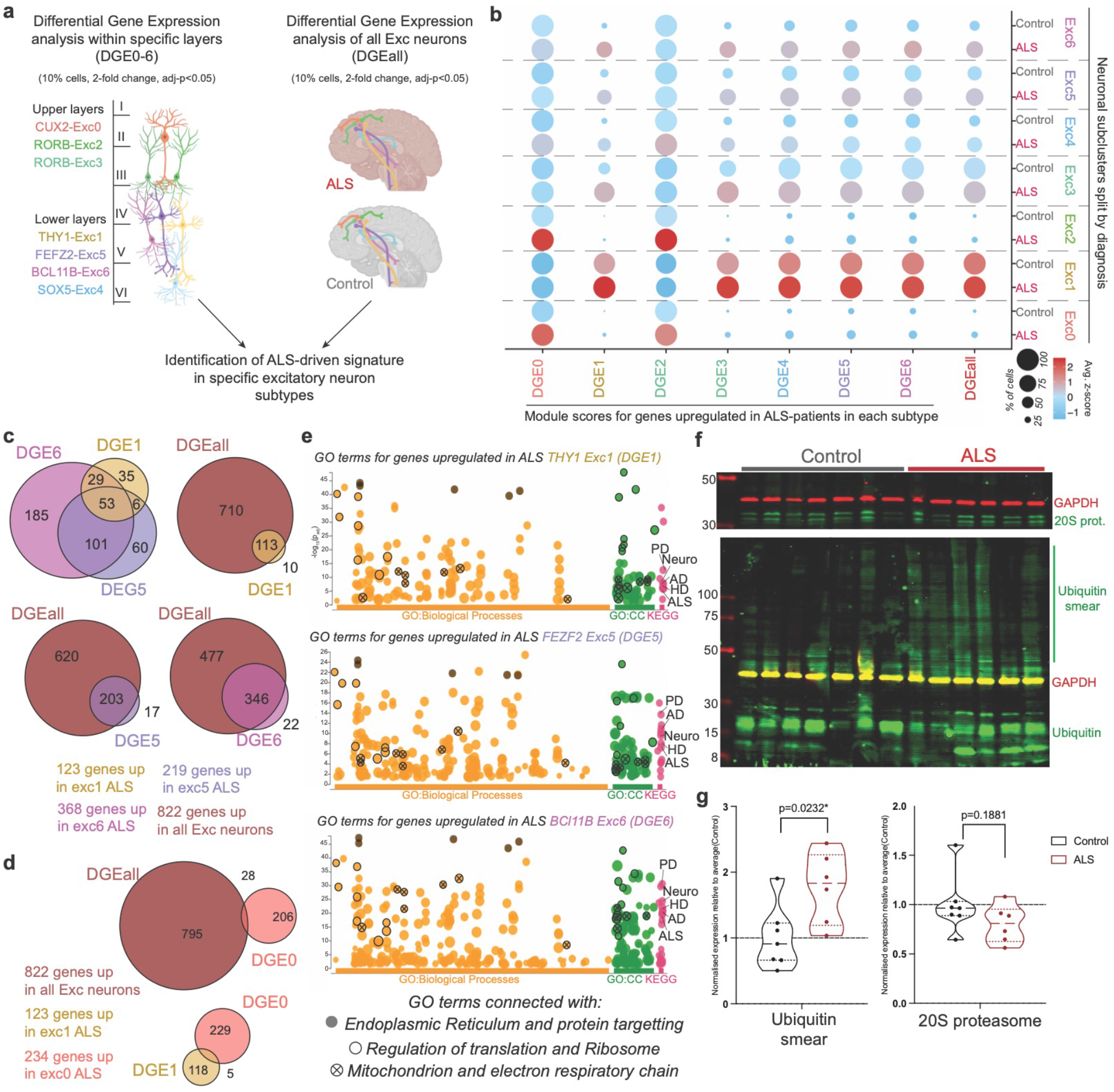
ALS excitatory neurons present increased expression of stress-related pathways. **a**, Schematic of Differential Gene Expression Analysis strategy. **b**, Dotplot representing the scores for genes upregulated in each subgroup of excitatory neurons (DGE0-6) and globally upregulated in all excitatory cells (DGEall). **c**, Comparison of genes globally upregulated in ALS (DGEall) with genes upregulated in specific layers CUX2-exc0 and THY1-exc1 (genes expressed by >10% of cells, >2-FC, adjusted p-value<0.05). **d**, Comparison of genes globally upregulated in ALS (DGEall) with genes upregulated in classes of L5-ETNs (genes expressed by >10% of cells, >2-FC, adjusted p-value<0.05). e, Gene Ontology analysis for genes upregulated in L5-ETNs classes (DGE1,5,6), highlighted terms are shared between the three and fall in categories described (CC=Cellular Components). **f-g**, Western Blot quantification of ubiquitin accumulation and 20S proteasome subunit from MotorCortex of ALS patients and age-matched controls.

Subsequent Gene Ontology (GO) analysis showed that DEGs in *CUX2-cells* were associated with synaptic biology (Extended Data Figure 4a,b), which could be due to changes in synaptic activity of degenerating neurons in deeper cortical layers. In contrast, DEGs identified in classes of ETNs were connected to cellular stresses previously associated with ALS^1,2^, even from studies with hundreds of patients^52^ (Fig. 3e). Interactome analysis confirmed coordinated alterations in the expression of genes that function in translational machinery, mitochondria, protein folding, proteostasis and degradation pathways connected to the proteasome and shared many transcriptional changes with patients’ excitatory cells as a whole (Extended Data Fig.4c,d-5). Interestingly, these pathways were specifically upregulated in neurons of deeper cortical layers rather than upper layer (Extended Data Fig. 4e,f). Comparison with other studies underlined similarities with genes upregulated in excitatory neurons from MS patients^18^ but not neurons from AD patients^20^ (Extended Data Fig. 4g,h), suggesting that similar processes might be at the base of neurodegeneration but these changes are not universal.

Presently, *in vitro* modelling of sporadic ALS requires high numbers of lines, high-throughput methods and needs further standardization^53–55^, we therefore decide to implement a system that would allow to probe disruptions of proteostasis in human neurons and test if any of these changes parallel the disruptions seen in ALS patients to interpret what proportion of this complex transcriptomic signature may be associated with proteostatic stress specifically in neuronal cells. In order to do so, we implemented transient proteasome inhibition as a model to induce TDP-43 nuclear loss as seen in patients’ Betz cells^49^, phase separation^56^, stress granules^57^ and other ALS-related dysfunction in human neurons^5,58^ (Extended Data Fig. 6a). To recapitulate proteostatic stress we applied a proteasome inhibitor to human Pluripotent Stem Cells (hPSC)-derived neurons^58,59^ and induced nuclear loss of TDP-43 (Extended Data Fig. 6b,c). Bulk RNA-sequencing analysis showed widespread changes after treatment, with a significant overlap of upregulated genes between stressed hPSC-neurons and sALS-neurons, specifically proteasome subunits and heat-shock response-associated chaperonins and GO analysis of shared genes confirmed the upregulation of proteasomal and chaperone complexes (Extended Data Fig. 6d-g). Moreover, genes upregulated in both conditions show a significant overlap with transcripts misregulated after downregulation of TDP-43 in neurons^58^ (Extended Data Fig. 6h). This confirms that some changes identified in sALS neurons are connected to neuronally intrinsic proteostatic alternations and at least in part connected to alterations in TDP-43.

To confirm hindered proteostasis in ALS cortex, we selected a second cohort of sALS patients and controls. We extracted protein, confirmed increased insoluble TDP-43 in patients (Extended Data Fig. 6i-j) and showed that, despite the presence of core proteosomal subunits, pathology is accompanied by the accumulation of highly ubiquitinated proteins (Fig. 3f,g), hallmark of impaired proteostasis. These findings suggest that proteasome inhibition orchestrate alterations like those observed in ETNs from ALS patients, underscoring the connection between neuronal stress and loss of proteostatic homeostasis.

### Oligodendroglial cells respond to neuronal stress with a neuronally-engaged state

To reach deep into the cord ETNs are dependent on robust axonal integrity^60^ and because others detected changes in myelination in ALS motor cortex^16^ and in FTD frontal cortex^27^, we analysed nuclei from myelinating cells. 19,151 nuclei from oligodendroglia were clustered in five groups: one of OPCs – Oliglia3, and four of oligodendrocytes – Oliglia0,1,2,4 (Fig. 4a-c, Extended Data Fig. 7a-b). We noted a significant depletion of ALS- nuclei in Oliglia0 whereas Oliglia1 and Oliglia4 were enriched in patients (Fig. 4d). GO analysis for genes enriched in each group revealed that Control-enriched Oliglia0 was characterised by terms connected to oligodendrocyte development and myelination and expressed higher levels of myelinating genes, e.g. *CNP, OPALIN, MAG* (Fig. 4e, Extended Data Fig. 7c-e). Conversely, ALS-enriched Oliglia1 showed terms for neurite morphogenesis, synaptic organization and higher expression of synaptic genes *DLG1, DLG2, GRID2* (Fig. 4f, Extended Data Fig. 7f-h). Intriguingly, expression of neuronal RNAs has been found in of oligodendrocytes in primate motor cortex^41^.

**Fig. 4.**
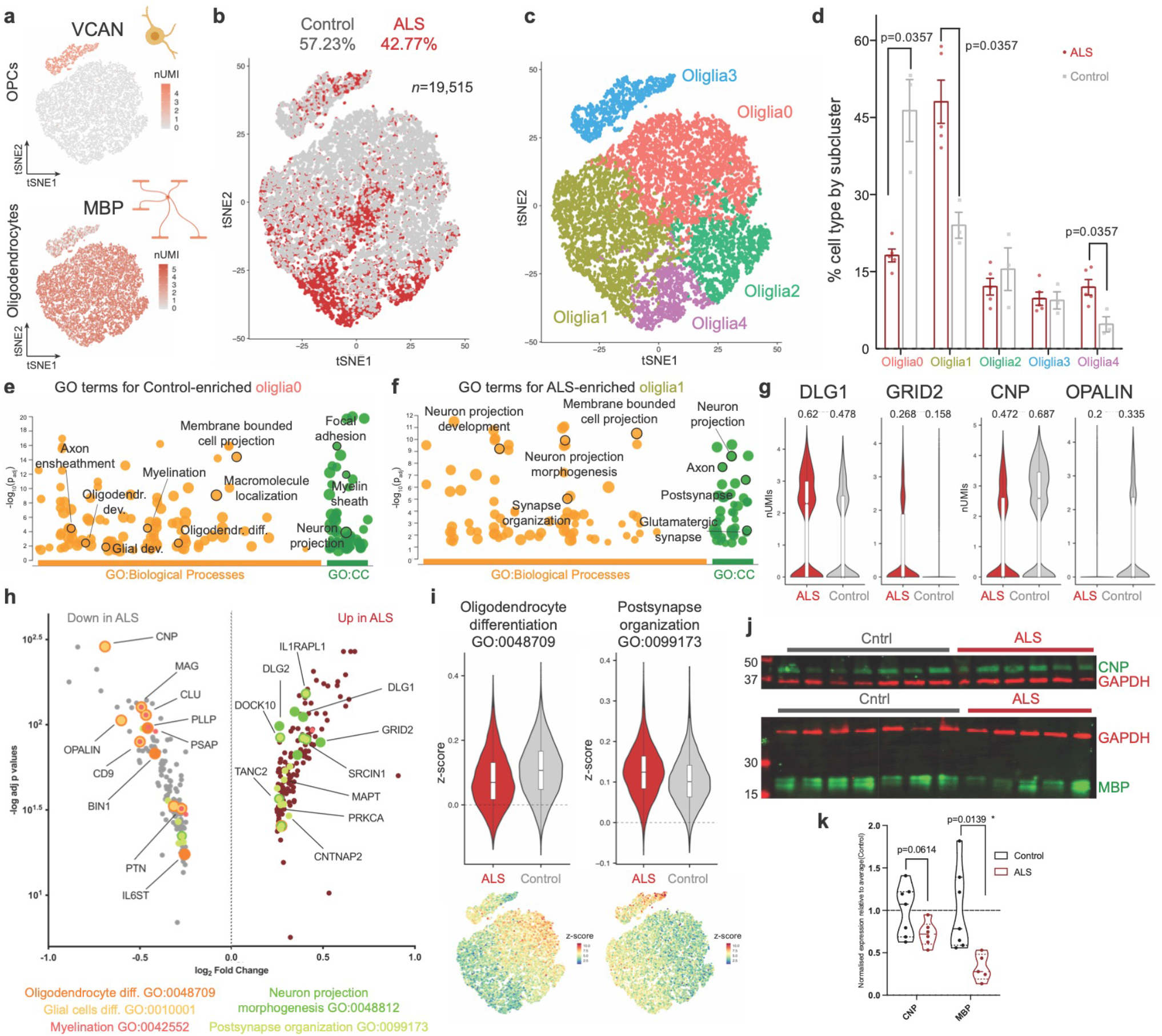
In ALS, oligodendroglial cells decrease their myelinating machinery in favour of a neuro-engaged state. **a**, *t*-SNE projection of markers of OPCs and oligodendrocytes. **b**, *t*-SNE projection of oligodendroglial cluster (ALS *n*=8,372 nuclei, Control *n*=11,168 nuclei). **c**, *t*-SNE projection of subclusters identified within oligodendroglia. **d**, Distribution of subclusters by diagnosis (U-Mann Whitney test). **e**, Gene Ontology analysis for genes characteristic of Control-enriched oliglia0, highlighted terms involved in myelination (CC=Cellular Components). **f**, Gene Ontology analysis for genes characteristic of ALS-enriched oliglia1, highlighted terms involved in neuro-supportive functions (CC=Cellular Components). **g**, Violin plots of representative genes for neuro-supportive functions (left) and myelination (right) (fraction of cell expressing). **h**, Volcano plot of differentially expressed genes in oligodendroglia. Highlighted genes identified in GO terms related to myelination (orange) and neuro-supportive functions (green). **i**, Violin plots representing z-score for selected GO terms and related *t*-SNE projection. **j-k**, Western Blot quantification of CNPase and MBP from MotorCortex of ALS patients and age-matched controls.

Global differential gene expression analysis supports a shift from a myelinating to a neuronally-engaged state with upregulation of genes involved in synapse modulation and decrease of master-regulators of myelination, as confirmed by GO analysis (Fig. 4g-i, Extended Data Fig. 7j,k). Loss of myelination is exemplified by the expression of G-protein coupled receptors (GPRCs) that mark developmental milestones: *GPR56*, expressed in OPCs^61^, and *GPR37*, expressed in myelinating cells^62^, were lowly expressed in ALS-enriched subgroups and globally downregulated (Extended Data Fig. 7i).

To further explore these changes, we compared them with published reports that identified shifts in oligodendrocytes (Extended Data Table 4)^19^. Comparison of Jäkel et al.^19^ with our study revealed that Control-enriched Oliglia0 most closely resembled highly myelinating, *OPALIN^+^* cells from Jäkel6 (Extended Data Fig. 8a,b), while ALS-enriched Oliglia1 and Oliglia4 aligned to not-actively myelinating Jäkel1 (Extended Data Fig. 8c-h). To confirm this shift, we ran validations on protein extracts from patients and controls and showed that oligodendrocyte-specific, myelin-associated proteins CNP and MBP are downregulated in motor cortices from patients (Fig. 4j-k), consistent with previous studies identifying demyelination in sALS patients^16^. The data so far shows how activation of stress pathways in deep layer neurons is accompanied by a shift in oligodendrocytes from active myelination to oligo-to-neuron contact (Extended Data Fig. 8i).

### Microglial activation is characterised by an ALS-specific endo-lysosomal response

Mouse models^63^, patient samples^6^ and function of ALS-related genes in myeloid cells^64–66^ have demonstrated the importance of microglia as modifiers of disease so we interrogated changes in this cell type. In the 1,452 nuclei from microglia (Fig. 5a, Extended Data Fig. 9a), we identified 159 genes upregulated in patients and, remarkably, many were associated with endocytosis and exocytosis, previously implicated in ALS^65,66^ (Fig. 5b). Several of these genes were also associated with microglial activation (*CTSD*) and neurodegenerative disorders (*APOE*) (Fig. 5c,d). Interestingly, genes associated with ALS/FTD were upregulated: *TREM2, OPTN, SQSTM1/p62, GRN* (Fig. 5e). GO analysis for upregulated genes confirmed a pro-inflammatory state highlighting activation of endo-lysosomal pathways, secretion and immune cell degranulation previously associated with myeloid cells in ALS^65,66^ (Fig. 5f,g). Further subclustering identified three groups: homeostatic Micro0, “Disease Associated Microglia”-like Micro1, and cycling Micro2 (Extended Data Fig. 9b,c). Notably, genes that characterised Micro1 were also upregulated in sALS (Extended Data Fig. 9d,e), in conjunction with a downregulation of homeostatic genes and upregulation of reactive pathways (Extended Data Fig. 9f-i).

**Fig. 5.**
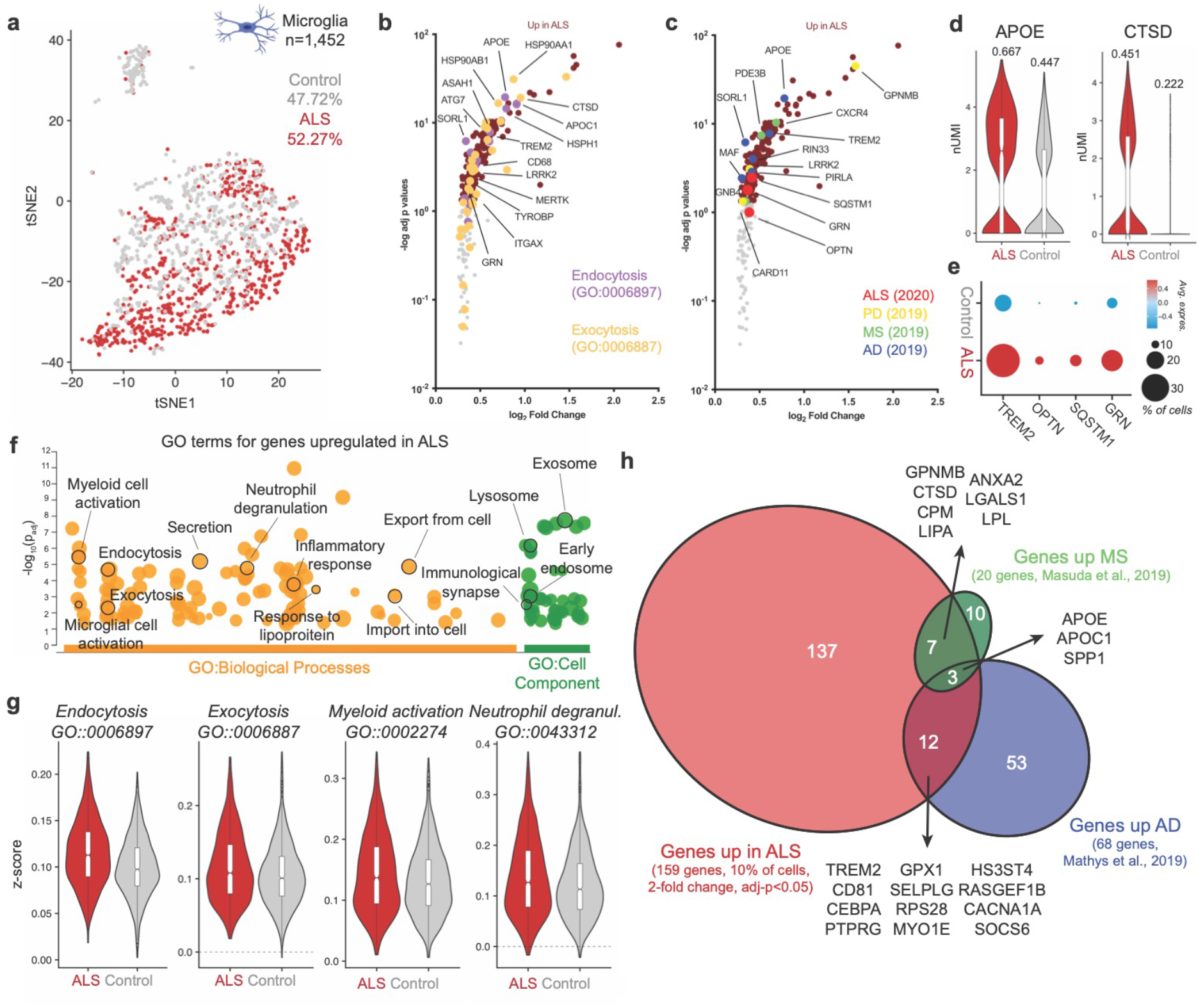
Disease-Associated Microglia signature in ALS. **a**, *t*-SNE projection of microglia (ALS *n*=759 nuclei, Control *n*=693 nuclei). **b,c**, Volcano plot of genes upregulated in microglia from ALS. Genes identified in Gene Ontology terms for endocytosis and exocytosis highlighted in b, genes associated to neurodegenerative diseases highlighted in c (ALS, PD - Parkinson’s disease, MS, AD). **d**, Violin plots of representative genes upregulated in ALS patients associated with reactive microglia (geometric boxplots represent median and interquantile ranges) (fraction of cell expressing). **e**, Dotplot representing expression of genes associated with ALS-FTD pathogenesis upregulated in microglia from patients. **f**, Gene Ontology analysis for genes upregulated in ALS microglia, highlighted terms involved in myeloid cells biology and/or pathogenesis of ALS. **g**, Violin plots representing z-score for selected, statistically significant GO terms from 4f. **h**, Comparison of genes upregulated in microglia from ALS patients with genes upregulated in microglia in other neurodegenerative diseases.

To identify modulators of this signature, we used the Connectivity Map (CMap) pipeline^67^, which contains gene expression data of 9 human cell lines after thousands of perturbations, allowing association between a given transcriptomic signature and a specific alteration. This analysis revealed that genes dysregulated in microglia positively correlated with regulators of cell cycle and senescence, suggesting an exhaustion of microglial proliferation. We also found a negative correlation with a type I-interferon-associated responses (*IFNB1*), which is targeted in treatments for neurological diseases to reduce inflammation^67^ (Extended Data Fig. 10a).

Given the stress signature identified in neurons, we wondered whether these transcriptomic changes might be driven by neuronal distress. We differentiated microglia-like cells (iMGLs)^68^ and neurons (piNs)^59^ from hPSCs, triggered neuronal apoptosis and then introduced apoptotic neurons to iMGLs *in vitro* (Extended Data Fig. 10b-c). Quantitative assessment of selected transcripts by RT-qPCR confirmed that dead neurons lead to significant downregulation of homeostatic genes (Extended Data Fig. 10d), upregulation of genes involved in the endo-lysosomal trafficking (*CTSD, ITGAX, LGALS3, SQSTM1/p62*) and downregulation of actively cycling cells markers (Extended Data Fig. 10e-f), suggesting that changes identified in microglia from patients are, at least in part, a response to neuronal apoptosis.

We next asked if these changes were a general response to neurodegeneration or restricted to ALS. By comparing our results with published snRNA-seq studies in AD^20^ and MS^69^, we identified dysregulation of lipid metabolism (*APOE, APOC1, SPP1*) as a common feature in microglia, and genes associated with DAMs shared between ALS and MS (*CTSD, GPNMB, CPM, LPL*) and ALS and AD (e.g. *TREM2*) (Fig. 5h). Genes specifically upregulated in ALS were related to vesicle trafficking, myeloid cell degranulation and the lysosome (e.g., *SQSTM1/p62, LGALS3, GRN, ASAH1, LRRK2*). This evidence suggests the induction of a shared microglial reactive state in neurodegenerative diseases, yet in ALS neuronal death activates changes connected to dysfunctional endo-lysosomal pathways.

## Discussion

A key question in the study of neurodegeneration is why certain cell types are more susceptible to different diseases^70^. In this study, we identified the enrichment for ALS/FTD associated genes in a class of ETNs which provides a connection between this neuronal subtype and its propensity to accumulate TDP-43 aggregates^11,13,50^ leading to their gradual loss in ALS/FTD^10^. This enrichment is not recapitulated for risk factors connected to AD and MS, related to immune processes and more enriched in microglia^30^. One study suggested that ALS-associated variants connected to autophagy and protein clearing are most highly expressed in glutamatergic neurons^51^ and these findings were later corroborated adding the importance of axonal dynamics and ribonucleotide metabolism^52^, here we provide a more detailed dissection of which subtype that might be.

Additionally, we identified a broadly shared transcriptomic signature of cellular stress pathways in classes of deep layer excitatory neurons. These alterations in RNA translation and proteostasis have previously been involved in models of ALS^1,2^. Our study recapitulates these changes, it highlights their cell-type specificity and links them to the identification of rare mutations in regulators of these pathways in familial forms of ALS^71^. These molecular mechanisms are confirmed to be connected to proteasomal function and protein homeostasis by our neuronal human *in vitro* model. The nuclear nature and the low-coverage of this kind of sequencing but also the small sample size in our study does not allow for further, confident dissection of the specifically neuronal changes in RNA biology identified in *in vitro* models and patient samples^58,72^. It remains intriguing to speculate how RNA metabolism and proteostasis might be mis-regulated in ETNs, mouse models where these pathways are specifically altered in CSMNs might shed a light on their interplay in this specific neuronal type.

We suggest two mechanisms by which ETNs are rendered more susceptible to ALS: 1. the intrinsically higher expression of risk factors; 2. processes of neurodegeneration in classes of ETNs that might exacerbate and contribute to vulnerability of these cells in a combinatorial effect. Recent snRNA-seq studies have unravelled susceptibility of specific neuronal types in other diseases: mid-layer *RORB6^+^* neurons in AD^22,73^; upper layer *CUX2-* neurons in MS^18^; dopaminergic neurons in Parkinson’s Disease^23^; ETNs affected in ALS/FTD as described by our study and spinal cord motor neurons as suggested in recent reports^74,75^. Impairment of proteostatic mechanisms seems to be a common theme in degenerating neurons regardless of the disease, however, only in ALS these changes are specifically connected to upregulation of transcripts connected to RNA metabolism, trend that appears to go in opposite direction in AD^73^. Integrative analyses of these studies might mark the beginning of a new era in the understanding of the mechanisms behind selective neuronal vulnerability to different diseases.

Emerging studies have shown that glial cells are important modifiers in ALS/FTD^16^. We show that changes in processes involved in oligodendrocyte differentiation and myelination may contribute to neuronal degeneration and/or be a coordinated response to ALS and appear to contrast those described in MS^19^. Moreover, we revealed perturbations in key myelin-regulators, such as *OPALIN, CNP*, and *MAG*, across oligodendrocyte clusters but in these cells only, as opposed to AD where myelination-related changes were present across multiple cell types^20,24,25^. Given the similarities in the stress signature identified in neurons with changes in MS lesions but not in AD, it is puzzling how changes in myelination might be a consequence or cause of neuronal degeneration.

Intriguingly, recent work showed expression of neuronal RNA in oligodendrocytes in human motor cortex^41^. Upregulation of synaptic transcripts in this cell type in ALS patients might represent phagocytic activity^76^ or the need for synaptic proteins for deposition of myelin sheath^77^. These speculations are interesting if coupled with the upregulation of synaptic machinery in upper layer *CUX2*-neurons and the documented loss of postsynaptic molecules in ETNs in ALS^78^. Moreover, snRNAseq studies of FTD cortices identified changes in myelinating cells in response to neuronal loss and underlined the importance of cell-to-cell communication^27^. Finally, recent GWAS studies have pointed at excitatory neurons, myelinating cells and inhibitory neurons as more sensitive to genetic risks for ALS^52^. These observations suggest a coordinated response of the whole Cortico-Spinal motor circuit in an attempt to compensate for loss of inputs to the cord. Further investigations could focus on shifting oligodendroglial states in disease models and determine changes during disease progression in the scope to complement efforts aimed to controlling neuronal activity^79^.

Finally, we found distinct perturbations in ALS-associated microglia, particularly in endo-lysosomal pathways. We and others have implicated ALS/FTD-associated gene *C9orf72* in endosomal trafficking and secretion in myeloid cells^65,66^ and the upregulation of lysosomal constituents, e.g. *CTSD*, was identified in this study and by others in patients^80^. Coupled with the upregulation of ALS/FTD-associated genes *SQSTM1/p62, OPTN, TREM2* and *GRN*, this suggests a mechanistic convergence on vesicle trafficking and inflammatory pathways that may initiate and/or exacerbate the homeostatic-to-DAM transition in ALS. The interferon-response-related changes we delineate, as identified by others in *C9orf72*-ALS^81^, provide a parallel between sporadic and familial ALS. Overall, these changes had partial overlap with those in microglia in AD^20,21^ and in MS^69^, suggesting that drugs specifically modulating myeloid cells in neurodegenerative diseases may provide a basis for new therapeutic approaches for ALS/FTD and warrants further study. Recent reports have shown how Disease Associated Microglia and their activity might actually be beneficial in disease contexts^82^, studies specifically manipulating microglial states might elucidate the “friend or foe” role of these cells in ALS^37^.

In summary, we show that classes of ETNs require the expression of a collection of genetic risk factors for ALS/FTD with pivotal roles in proteostasis. This intrinsically higher expression of disease-associated genes might be at the bottom of a “first over the line” mechanism leading to disruption of homeostasis in groups of deep-layer excitatory neurons. These alterations trigger a cascade of responses: superficial neurons upregulate synaptic genes; oligodendroglia shift from a myelinating to a neuronally-engaged state; microglia activate a pro-inflammatory signature. Our study offers a view in which neurocentric disease vulnerability sparks responses in other neuronal types and glial cells, but it also shows that enrichment of ALS/FTD-related genes in ETNs is also coupled with processes engaging disease related genes in different cells, i.e. microglia. This view is a first insight into the disruptions of cortical biology in ALS and provides a connection between changes in cellular components and mechanisms associated with ALS^83^. Future investigations should consider multicellular disruptions in ALS/FTD, where the survival of the neuron is unmistakably pivotal, but targeting other cells to reduce inflammation, promote myelination, and bolster neuronal circuitry may re-establish a neuroprotective environment.

## Limitations of this study

One limitation of this study is the small size of the cohort. ALS is a very heterogenous disease^7^, and smaller cohort sizes cannot fully recapitulate its etiological diversity. However, only recently biobanks have been able collect enough samples to generate reports with dozens individuals^7,8^, we hope that the increase in samples availability and affordability of single-cell technology will allow a more comprehensive view of transcriptomic changes in ALS. A bigger cohort size would also allow a more stringent analysis of differentially expressed genes (such as by “pseudo-bulking”) before nominating DEGs. We also recognise that our study would benefit from additional validation at RNA and/or protein level, we hope that in the future this kind of studies would accompany bioinformatics analyses with more validations. This would also elucidate some of the findings in this study. For example, are oligodendrocytes in ALS patients really expressing higher levels of neuronal genes or is this an artifact coming from contaminations^84^? Nonetheless, we believe that this study provides original and novel insights the involvement of different cell types in ALS and a different view in the motor cortex of ALS patients.

## Methods

### Human donor tissue

Frozen post-mortem human cortical samples from cases of sporadic ALS patients and age-matched controls were obtained from the Target ALS Neuropathology Core that drew upon the repositories of five institutions. Specimens from the medial, lateral, or unspecified motor cortex were grouped together. Additional post-mortem human samples of the posterior frontal cortex consistent with the motor cortex from ALS patients and controls were obtained at MGH using a Partners IRB approved protocol and stored at −80°C.

### Isolation of nuclei

RNA quality of brain samples was assessed by running bulk nuclear RNA on an Agilent TapeStation for RIN scores. Extraction of nuclei from frozen samples was performed as previously described^85^. Briefly, tissue was dissected and minced with a razor blade on ice and then placed in 4 ml ice-cold extraction buffer (Wash buffer (82 mM Na2SO4, 30 mM K2SO4, 5 mM MgCl2, 10 mM glucose, and 10 mM HEPES, pH adjusted to 7.4 with NaOH) containing 1% Triton X-100 and 5% Kollidon VA64). Tissue was homogenized with repeated pipetting, followed by passing the homogenized suspension twice through a 26 ½ gauge needle on a 3 ml syringe (pre-chilled), once through a 20 mm mesh filter, and once through a 5 mm filter using vacuum. The nuclei were then diluted in 50 ml ice-cold wash buffer, split across four 50 ml tubes, and centrifuged at 500xg for 10 minutes at 4°C. The supernatant was discarded, the nuclei pellet was resuspended in 1 ml cold wash buffer.

### 10X loading and library preparation

Nuclei were DAPI-stained with Hoechst, loaded onto a hemocytometer, and counted using brightfield and fluorescence microscopy. The solution was diluted to ~176 nuclei/ul before proceeding with Drop-seq as described in ref.15^28^. cDNA amplification was performed using around 6000 beads per reaction with 16 PCR cycles. The integrity of both the cDNA and tagmented libraries were assessed for quality control on the Agilent Bioanalyzer as in ref^86^. Libraries were sequenced on a Nova-seq S2, with a 60 bp genomic read. Reads were aligned to the human genome assembly (hg19). Digital Gene Expression files were generated with the Zamboni Drop-seq analysis pipeline, designed by the McCarroll group^85^,^87^.

### Filtering of expression matrices and clustering of single nuclei

A single matrix for all samples was built by filtering any barcode with less than 400 genes and resulting in a matrix of 27,600 genes across 119,510 barcodes. This combined UMI matrix was used for downstream analysis using Seurat (v3.0.2)^29^. A Seurat object was created from this matrix by setting up a first filter of min.cells=20 per genes. After that, barcodes were further filtered by number of genes detected nFeature_RNA>600 and nFeature_RNA<6000. Distribution of genes and UMIs were used as parameters for filtering barcodes. The matrix was then processed via the Seurat pipeline: log-normalized by a factor of 10,000, followed by regressing out UMI counts (nCount_RNA), scaled for gene expression.

After quality filtering, 79,830 barcodes and 27,600 genes were used to compute SNN graphs and *t*-SNE projections using the first 10 statistically significant Principal Components. As previously described^88,89^, SNN-graphed *t*-SNE projection was used to determine minimum number of clusters obtain at resolution=0.2 (FindClusters). Broad cellular identities were assigned to groups on the basis of differentially expressed genes as calculated by Wilcoxon rank sum test in FindAllMarkers(min.pct=0.25, logfc.threshold=0.25). One subcluster with specifically high ratio of UMIs/genes was filtered out resulting in 79,169 barcodes grouped in 7 major cell types of the CNS: excitatory neurons, oligodendrocytes, inhibitory neurons, astrocytes, endothelial cells, microglia, oligodendrocyte progenitor cells (OPCs). Markers for specific cell types were identified in previously published human scRNAseq studies^30,31^.

Analysis of cellular subtypes were conducted by subsetting each group. Isolated barcodes were re-normalised and scaled and relevant PCs were used for re-clustering as a separate analysis. This newly scaled matrix was used for Differential Gene Expression analysis with the MAST algorithm^90^ in Seurat R package as previously reported^19,21,23,25,26^ with parameters FindAllMarkers(min.pct=0.10, logfc.threshold=0.25) and subclustering for identification of subgroups. Gene scores for different cellular subclusters were computed in each re-normalised, re-scaled sub-matrix using the AddModuleScore function in Seurat v3.0.2.

Re-analysis of publicly available datasets was performed using matrices and metadata available. Only barcodes with available metadata concerning their cellular identity were selected to use identities assigned by peer review publication^18,48^. The available barcodes were then loaded into Seurat v4.0.1^91^. Gene scores for different cellular subclusters were computed in each re-normalised, re-scaled sub-matrix using the AddModuleScore function as previously. Re-analysis of spatial transcriptomic from et al. was performed using publicly available data and codes from publication itself^47^.

### Gene Ontology, Interactome and Gene Set Enrichment Analyses

For GO terms analysis, we selected statistically significant up-regulated or down-regulated genes identified in each subcluster as described before (adj p-values<0.05, LFC=2). These lists were fed in the gProfiler pipeline^92^ with settings: use only annotated genes, g:SCS threshold of 0.05, GO cellular components and GO biological processes (26^th^ of May 2020 – 9^th^ of December 2021), only statistically significant pathways are highlighted. For oligodendrocytes cells (Extended Data Fig.8) statistically significant up-regulated genes identified in each subcluster as described before (adj p-values<0.05, LFC=2) were used for synaptic specific Gene Ontology analysis using SynGO^93^ (12^th^ of June 2020). Interactome map was built using STRING^94^ protein-protein interaction networks, all statistically significant upregulated genes were used, 810 were identified as interacting partners using “experiments” as interaction sources and a medium confidence threshold (0.400), only interacting partners are shown in Extended Data Figure 6. Gene Set Enrichment Analysis was performed using GSEA software designed by UC San Diego and the Broad Institute (v4.0.3)^95^. Briefly, gene expression matrices were generated in which for each subcluster each individual was a metacell, lists for disease-associated risk genes were compiled using available datasets (PubMed – ALS/FTD – Supplementary Table 2) or recently published GWAS for AD^33,34^ and MS^35^.

### Generation of Microglia-like Cells

Microglial-like cells were differentiated as described^68^ with minor modifications^37,88^. Briefly, hPSCs were cultured in E8 medium (Stemcell technologies) on Matrigel (Corning), dissociated with Accutase (Stemcell technologies), centrifuged at 300xg for 5 minutes, resuspended in E8 medium with 10*μ*M Y-27632 ROCK Inhibitor, 2M cells are transferred to a low-attachment T25 flask in 4ml of medium and left in suspension for 24 hours. The first 10 days of differentiation are carried out in iHPC medium: IMDM (50%, Stemcell technologies), F12 (50%, Stemcell technologies), ITSG-X 2% v/v (ThermoFisher), L-ascorbic acid 2-Phosphate (64 ug/ml, Sigma), monothioglycerol (400 mM, Sigma), PVA (10 mg/ml; Sigma), Glutamax (1X, Stemcell technologies), chemically-defined lipid concentrate (1X, Stemcell technologies), non-essential amino acids (NEAA, Stemcell technologies). After 24h (day0), cells are collected and differentiation is started in iHPC medium supplemented with FGF2 (Peprotech, 50 ng/ml), BMP4 (Peprotech, 50 ng/ml), Activin-A (Peprotech, 12.5 ng/ml), Y-27632 ROCK Inhibitor (1 *μ*M) and LiCl (2mM) and transferred in hypoxic incubator (20% O_2_, 5% CO_2_, 37°C). On day 2, medium is changed to iHPC medium plus FGF2 (Peprotech, 50 ng/ml) and VEGF (Peprotech, 50 ng/ml) and returned to hypoxic conditions. On day4, cells are resuspended in iHPC medium supplemented with FGF2 (Peprotech, 50 ng/ml), VEGF (Peprotech, 50 ng/ml), TPO (Peprotech, 50 ng/ml), SCF (Peprotech, 10 ng/ml), IL-6 (Peprotech, 50 ng/ml), and IL-3 (Peprotech, 10 ng/ml) and placed into a normoxic incubator (20% O_2_, 5% CO_2_, 37°C). Expansion of haematopoietic progenitors is continued by supplementing the flasks with 1ml of iHPC medium with small molecules every two days. On day10, cells are collected and filtered through a 40mm filter. The single cell suspension is counted and plated at 500,00 cells/well of a 6 well plate coated with Matrigel (Corning) in Microglia differentiation medium: DMEM/F12 (Stemcell technologies), ITS-G 2%v/v (Thermo Fisher Scientific), B27 (2%v/v, Stemcell technologies), N2 (0.5%v/v, Stemcell technologies), monothioglycerol (200 mM, Sigma), Glutamax (1X, Stemcell technologies), NEAA (1X, Stemcell technologies), supplemented with M-CSF (25 ng/ml, Peprotech), IL-34 (100 ng/ml, Peprotech), and TGFb-1 (50 ng/ml, Peprotech). Induced Microglia-like cells (iMGLs) are kept in this medium for 20 days with change three times a week. On day 30, cells are collected and plated on poly-D-lysine/laminin coated dishes in Microglia differentiation medium supplemented with CD200 (100 ng/ml, Novoprotein) and CX3CL1 (100 ng/ml, PeproTech), M-CSF (25 ng/ml, PeproTech), IL-34 (100 ng/ml, PeproTech), and TGFb-1 (50 ng/ml, PeproTech) until day 40.

### Feeding of apoptotic neurons to Microglia-like Cells

For feeding assays, neurons were generated from human iPSCs using an NGN2 overexpression system as described previously^59,96,97^. Day30 hiPSC-neurons “piNs” were treated with 2*μ*M H_2_O_2_ for 24 hours to induce apoptosis. Apoptotic neurons were gently collected from the plate and the medium containing the apoptotic bodies was transferred into wells containing day40 iMGLs. After 24 hours, iMGLs subjected to apoptotic neurons and controls were collected for RNA extraction.

### RNA extraction and RT-qPCR analysis

RNA was extracted with the miRNeasy Mini Kit (Qiagen, 217004). cDNA was produced with iScript kit (BioRad) using 50 ng of RNA. RT-qPCR reactions were performed in triplicates using 20 ng of cDNA with SYBR Green (BioRad) and were run on a CFX96 Touch™ PCR Machine for 39 cycles at: 95°C for 15s, 60°C for 30s, 55°C for 30s.

### Generation of hiPSC-derived neurons for bulk RNA sequencing

Human embryonic stem cells were cultured in mTESR (Stemcell technologies) on matrigel (Corning). Neurons were generated from HuES-3-Hb9:GFP based on the motor neuron differentiation protocol previously described^58,98^. Upon completion of the differentiation protocol, cells were sorted via flow-cytometry based on GFP signal intensity to yield GFP-positive neurons that were plated on PDL/laminin-coated plates (Sigma, Life technologies). Neurons were maintained in Neurobasal medium (Life Technologies) supplemented with N2 (Stemcell technologies), B27 (Life technologies), glutamax (Life technologies), non-essential amino acids (Life technologies), and neurotrophic factors (BDNF, GDNF, CNTF), and were grown for 28 days before the application of the proteasome inhibitors MG132 for 48 hrs.

RNA was extracted using RNeasy Plus kit (Qiagen), libraries were prepared using the Illumina TruSeq RNA kit v2 according to the manufacturer’s directions, and sequenced at the Broad Institute core with samples randomly assigned between two flow chambers. The total population RNA-seq FASTQ data was aligned against ENSEMBL human reference genome (build GRCh37/hg19) using STAR (v.2.4.0). Cufflinks (v.2.2.1) was used to derive normalized gene expression in fragments per kilo base per million (FPKM). The read counts were obtained from the aligned BAM-files in R using Rsubread. Differential gene expression was analyzed from the read counts in DESeq2 using a Wald’s test for the treatment dosage and controlling for the sequencing flow cell.

### Western blot analysis

As previously described tissue was minced, lysed in RIPA buffer with protease inhibitors (Roche) and sonicated^99^. After centrifugation, the supernatant was collected as soluble fraction and the insoluble pellet was resuspended in 8M urea buffer (Bio-Rad, 1632103). After protein quantification by BCA assay (ThermoFisher), ten micrograms of proteins were preheated in Laemmli’s buffer (BioRad), loaded in 4-20% mini-PROTEAN® TGX™precast protein gels (BioRad) and gels were transferred to a PDVF membrane. Membranes were blocked in Odyssey Blocking Buffer (Li-Cor) and incubated overnight at 4°C with primary antibodies. After washing with TBS-T, membranes were incubated with IRDye® secondary antibodies (Li-Cor) for one hour and imaged with Odyssey® CLx imaging system (Li-Cor). Primary antibodies: TDP-43 (Peprotech 10782-2-AP); GAPDH (Millipore Cat# MAB374; CST 2118 (14C10)); MBP (ThermoFisher PA-1-10008); CNP (Abcam ab6319(11-5b)); 20S (Enzo BML-PW8195-0025); Ubiquitin (CST 3936T (P4D1)).

### Immunofluorescence assays

Cells were washed once with PBS, fixed with 4% PFA for 20 minutes, washed again in PBS and blocked for one hour in 0.1% Triton in PBS with 10% donkey serum. Fixed cells were then washed and incubated overnight with primary antibodies at 4°C. Primary antibody solution was washed and cells were subsequently incubated with secondary antibodies (1:2000, Alexa Fluor, Life Technologies) at room temperature for 1 hour, washed with PBS and stained with DAPI. Primary antibodies used: Tuj1 (R&D, MAB1195), TDP-43 (Peprotech 10782-2-AP). Images were analysed using FIJI.

### Proteasome activity assay

Neurons were sorted in 96-wells plates and, after two weeks of maturation, treated for 24 hours. Cells were washed with 1xPBS, exposed to ProteasomeGlo® (Promega, G8660) and incubated for 30 minutes at RT. Fluorescence was measured using a Cytation™3 reader (BioTek).

## Acknowledgements

We thank the study participants and staff at Massachusetts Alzheimer’s Disease Research Center for sequencing (NIA P50 AG005134). We thank the study participants and Kathleen Wilsbach, Lyle Ostrow and staff at Target ALS Neuropathology core and associated institutions for validation cohort samples. We thank UCB Pharma for partially funding these studies. D.M. acknowledges support from Massachusetts ADRC (5P50AG005134) pilot project and the NINDS (K08NS104270). We would also like to thank Dr Paul Tesar and his group for invaluable discussions on oligodendroglial biology. Some figures created with BioRender.com.

## Author contributions

The study was designed by F.L., D.M. and directed and coordinated by K.E. and S.A.M with input from B.S. and I.K. Manuscript writing by F.L. and D.M., with support from O.P and A.B. F.L. performed bioinformatics analysis with the help of S.D.G., D.M. and D.M. D.M. and I.C. supported obtaining post-mortem samples and carried out nuclei isolation and RNA-sequencing with M.G. and L.B.; M.T., O.P., A.B., A.C. and B.J.J. performed bioinformatics analyses of bulk RNA-sequencing and helped with protein and RNA validation with cellular models; J.M.M. performed analysis of published datasets; M.T. and B.S. contributed to microglial biology section; K.E. acquired primary funding.

## Competing interests

I.K. is an employee at UCB Pharma and holds stock options. K.E. is a cofounder of Q-State Biosciences, Quralis, Enclear Therapies and is group vice-president at BioMarin Pharmaceutical. K.E. and F.L. are authors on a pending patent “Single-nuclei characterization of amyotrophic lateral sclerosis frontal cortex” (US 2022,17535070).

## Additional information

We worked to ensure diversity in experimental samples through the selection of genomic datasets. One or more of the authors self-identifies as an underrepresented ethnic minority in science. One or more of the authors of this paper self-identifies as a member of the LGBTQ+ community.

---

**Extended Data Fig. 1.**
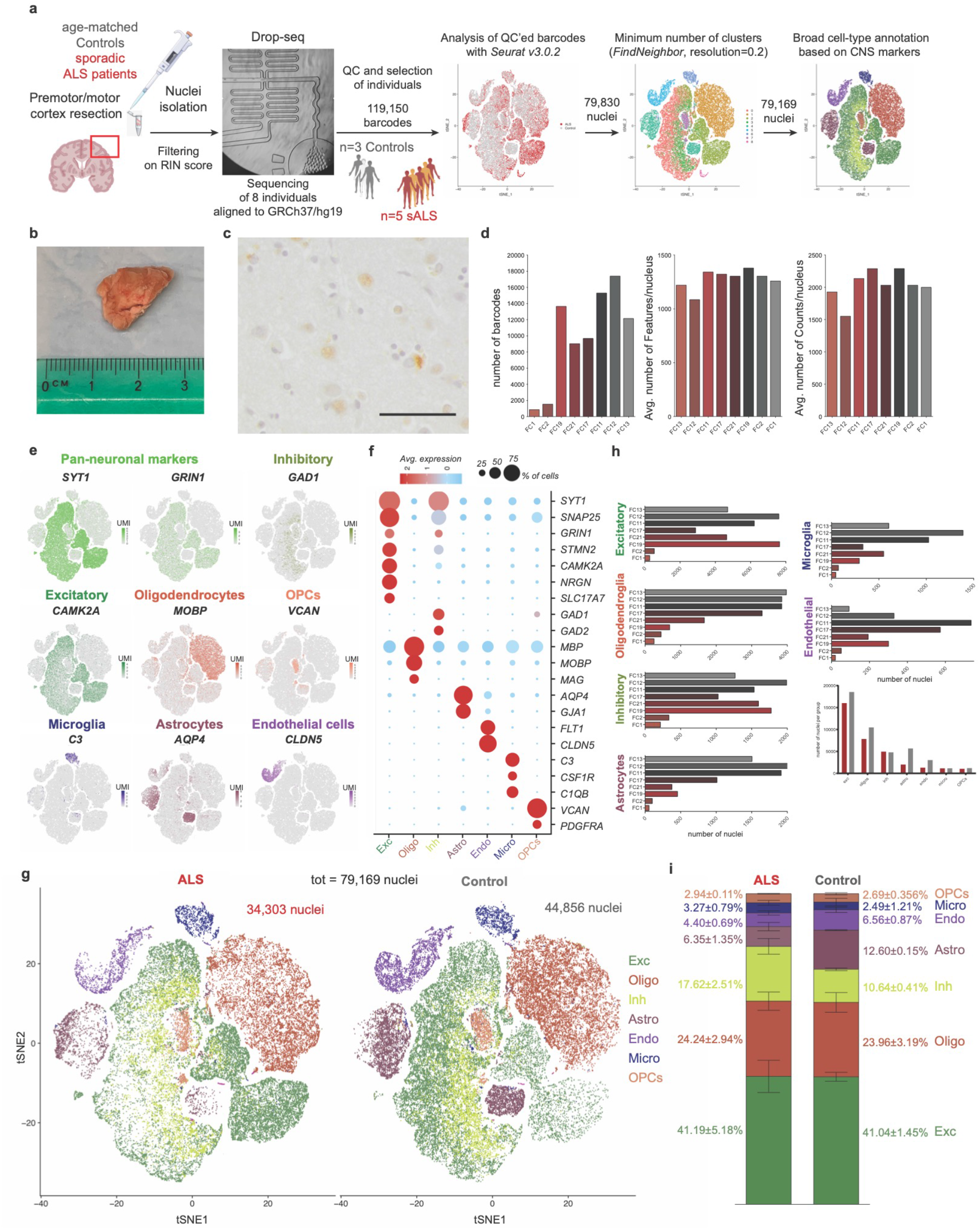
Technical parameters of snRNAseq and cell-type distribution across individuals. **a.** Schematic diagram of workflow for isolation of nuclei from cortices of ALS patients and age-matched controls followed by single-cell RNA sequencing by DropSeq, library generation and Quality Controls for analysis with Seurat 3.0.2 **b.** Frozen tissue from one of the individuals. **c.** Staining for TDP-43 in one of the patient sample, note neurons with skein-like inclusions and faint nuclear staining (scale bar 25*μ*m). **d.** Quality controls post-filtering (FC – Frontal Cortex): number of total nuclei detected (barcodes), average number of genes per nucleus (nFeatures), and average number of UMIs (Unique Molecular Identifiers) per nucleus (nCounts). **e.** *t*-SNE projections of the whole cohort with expression of broad cell type markers. **f.** Dotplot representing percentage of cells expressing additional cell type specific markers. **h**. *t*-SNE distribution of whole cohort with annotated cell types split by diagnosis (ALS patients *n*=5, age-matched Controls *n*=3, *n* = 79,169 total nuclei). **i.** Fraction of each cell types identified in whole cohort split by diagnosis.

**Extended Data Fig. 2.**
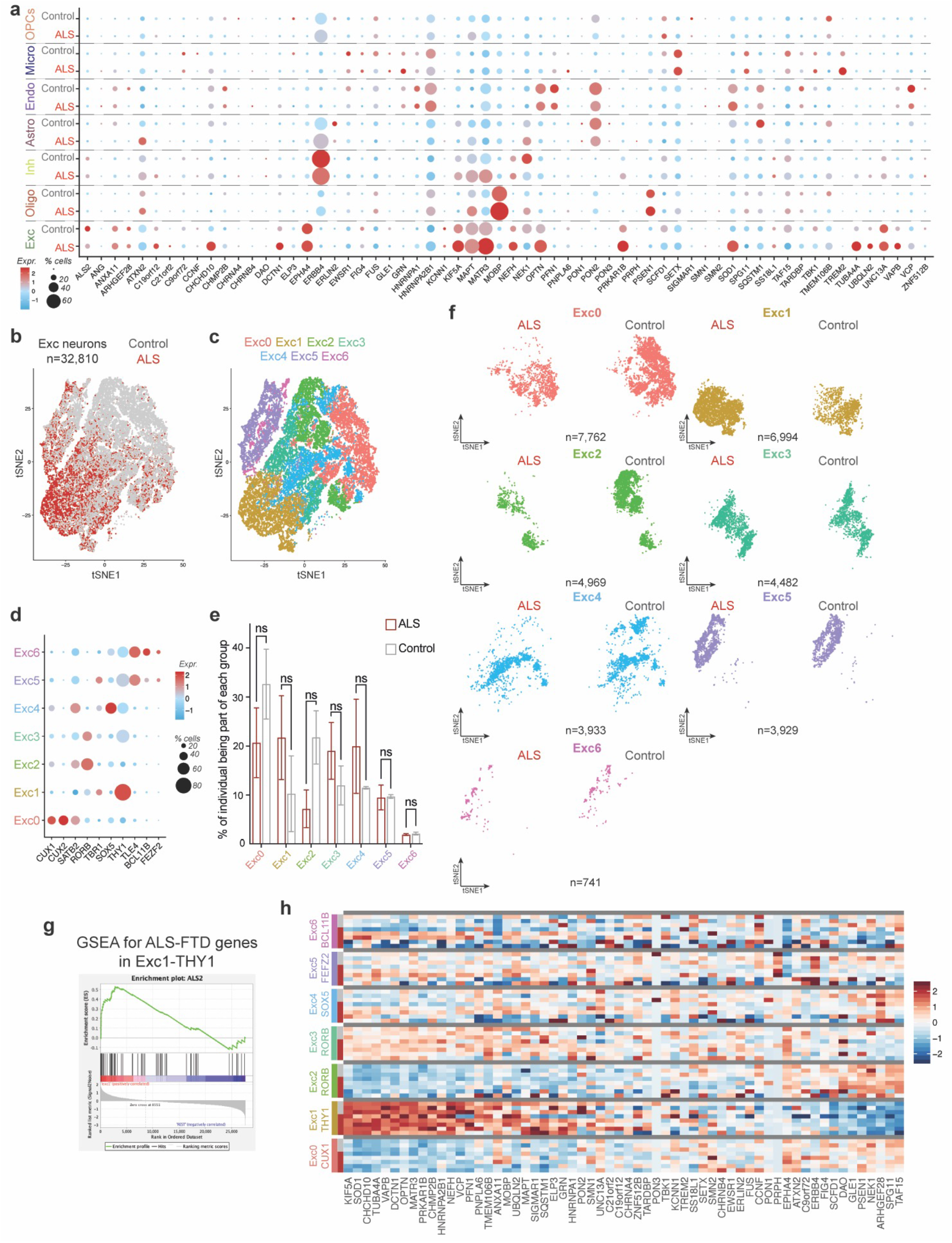
Expression of ALS-FTD associated genes in different cellular subtypes and excitatory neurons subtypes. **a.** Dotplot representing expression of gene associated with the ALS-FTD spectrum in each cell type identified in the whole cortex split by diagnosis. **b.** *t*-SNE projection of excitatory neurons clusters (ALS *n*=15,227 nuclei, Control *n*=17,583 nuclei). **c.** *t*-SNE projection of subclusters identified in excitatory neurons represents different, biologically relevant neuronal layers (*FindNeighbor*(res=0.2)). **d.** Dotplot representing percentage of cells expressing broad markers for different cortical layers. **e.** Distribution of excitatory neurons in subclusters by diagnosis. **f.** *t*-SNE projection of excitatory neurons by clusters and by diagnosis. **g.** Get Set Enrichment Analysis for the ALS-FTD associated genes in Exc1 cortical neurons. **h.** Heatmap representing expression of gene associated with the ALS-FTD spectrum in each excitatory neurons identified split by diagnosis.

**Extended Data Fig. 3.**
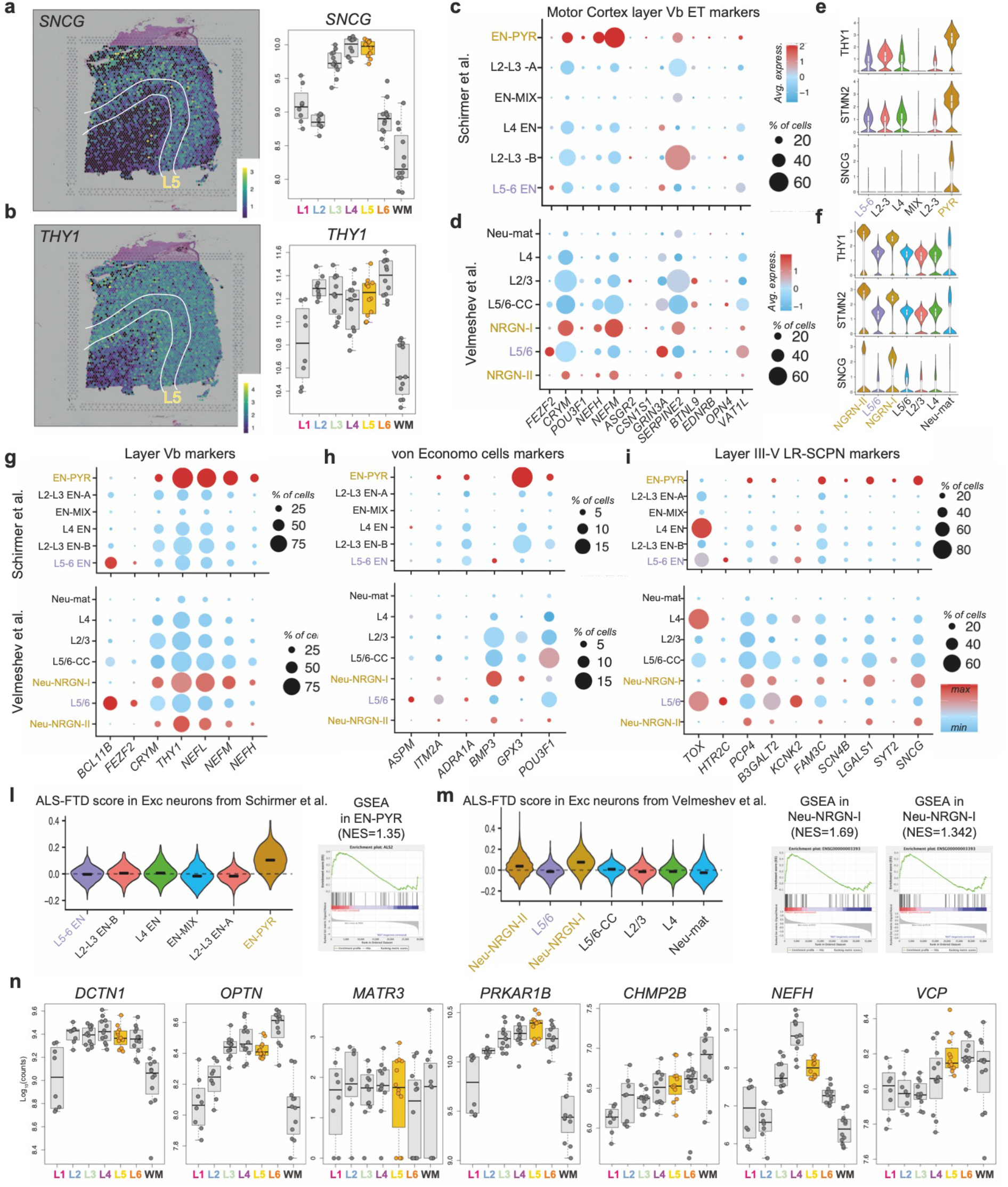
L5-ETNs/CSMNs-like neurons express higher levels of ALS-FTD related genes. **a,b**, Spotplot and corresponding boxplot from Maynard et al. for the expression of layer Vb Motor Cortex marker, *SNCG* and *THY1*, identified as enriched in Exc1. **c,d.** Dotplot and representative Violin plots for markers of L5 ExtraTelencephalic neurons of human Motor Cortex in Schirmer et al. **e,f.** Dotplot and representative Violin plots for markers of L5 ExtraTelencephalic neurons of human Motor Cortex in Velmeshev et al. **g-i.** Dotplot representing expression of Layer V markers (d), von Economo markers (e), LR-SCPN markers (f) in Schirmer et al. and Velmeshev et al. **l.** Violin plots and corresponding Gene Set Enrichment Analysis of z-scores for expression of ALS-FTD-associated genes in THY1-neurons identified by Schimer et al. (bars denote median). **m.** Violin plots and corresponding Gene Set Enrichment Analysis of z-scores for expression of ALS-FTD-associated genes in THY1-neurons identified by Velmeshev et al. (bars denote median). **n.** Boxplot from Maynard et al. for the expression of top 10 ALS/FTD associated genes identified in Exc1.

**Extended Data Fig. 4.**
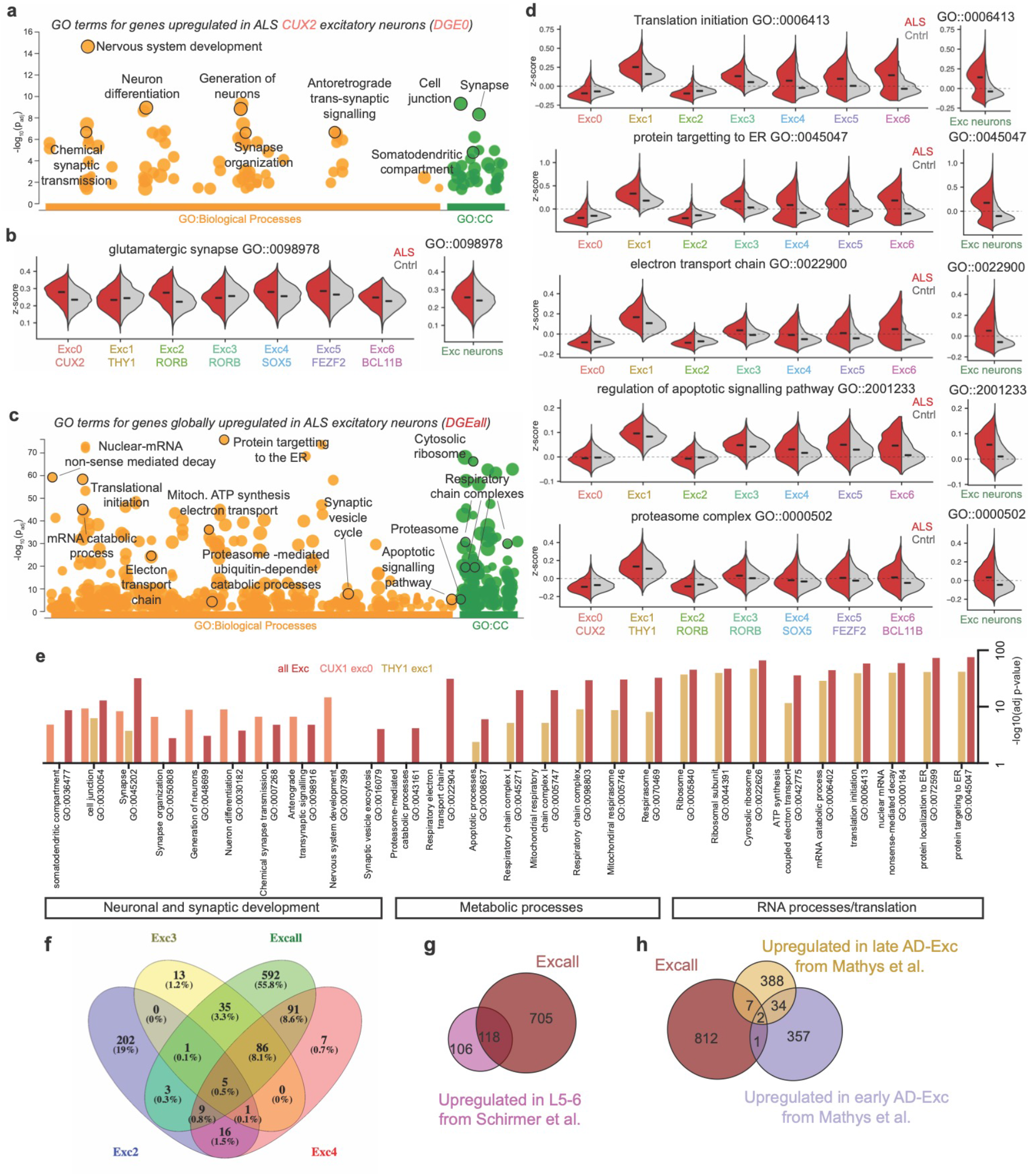
Classes of L5-ETNs express higher levels of stress pathways. **a.** Gene Ontology analysis for genes globally upregulated in ALS excitatory neurons (DGEall). **b.** Violin plots representing z-scores for selected, statistically significant GO terms upregulated in lower layers in each subgroup (left) and globally (right). **c.** Gene Ontology analysis of terms for genes upregulated in CUX2-Exc0 group (DGE0), highlighted terms involved in synaptic biology (CC=Cellular Components). **d.** Violin plot representing z-scores for selected, statistically significant GO terms upregulated in upper layers in each subgroup (left) and globally (right). **e.** Representation of -log10(adjusted p-values) of selected GO terms from previous figures for CUX2-Exc0 and THY1-Exc1 groups and globally. **f.** Venn Diagram depicting shared upregulated genes between other excitatory neurons and global signature. **g-h.** Venn Diagram depicting shared upregulated genes between treated MS excitatory neurons from Schirmer et al. and AD excitatory neurons from Mathys et al. with changes found in this study.

**Extended Data Fig. 5.**
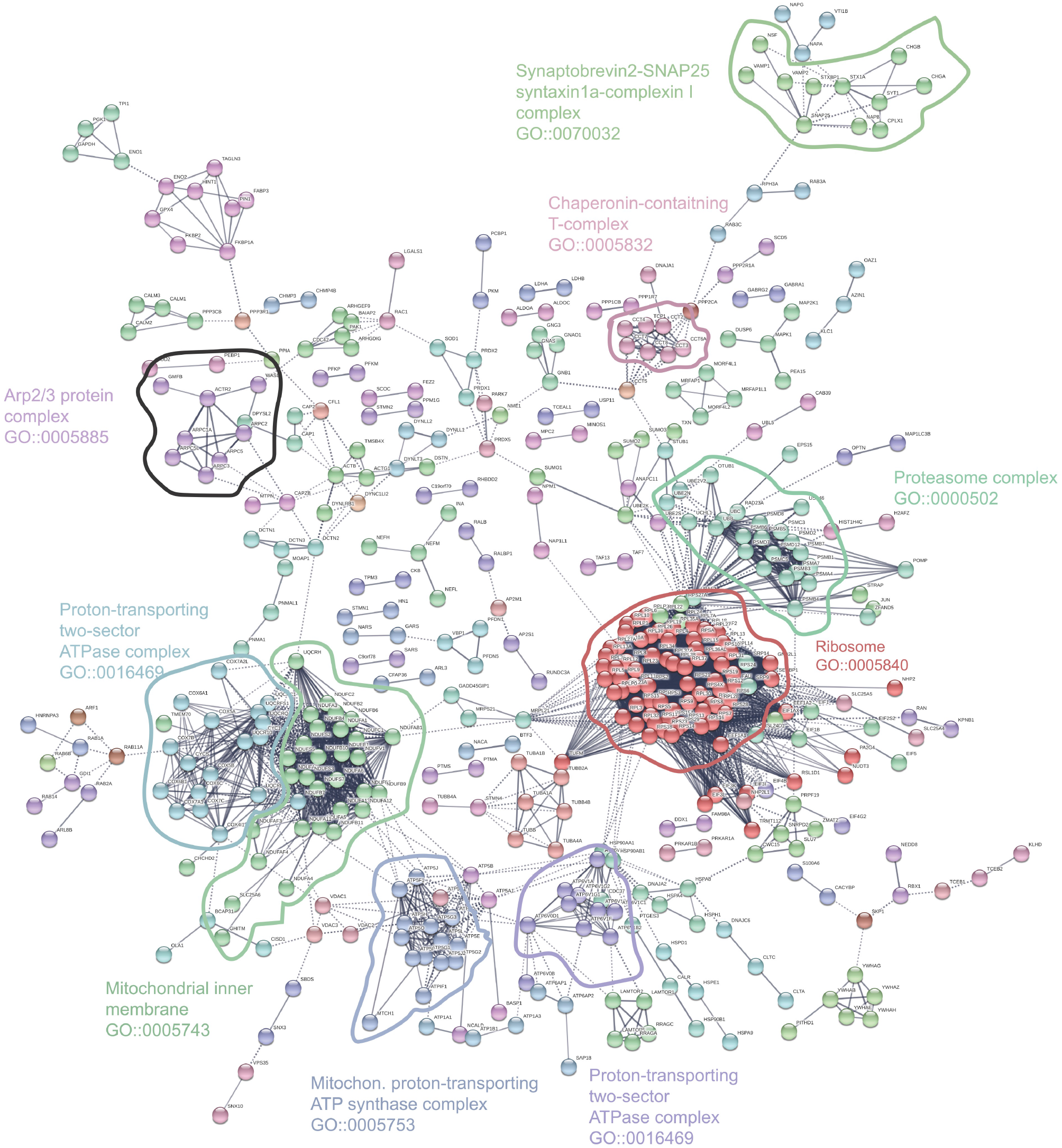
Global protein-protein interaction network for genes upregulated in ALS excitatory neurons. Color-coding derived from MLC clustering (4) to identified closely related groups of proteins.

**Extended Data Fig. 6.**
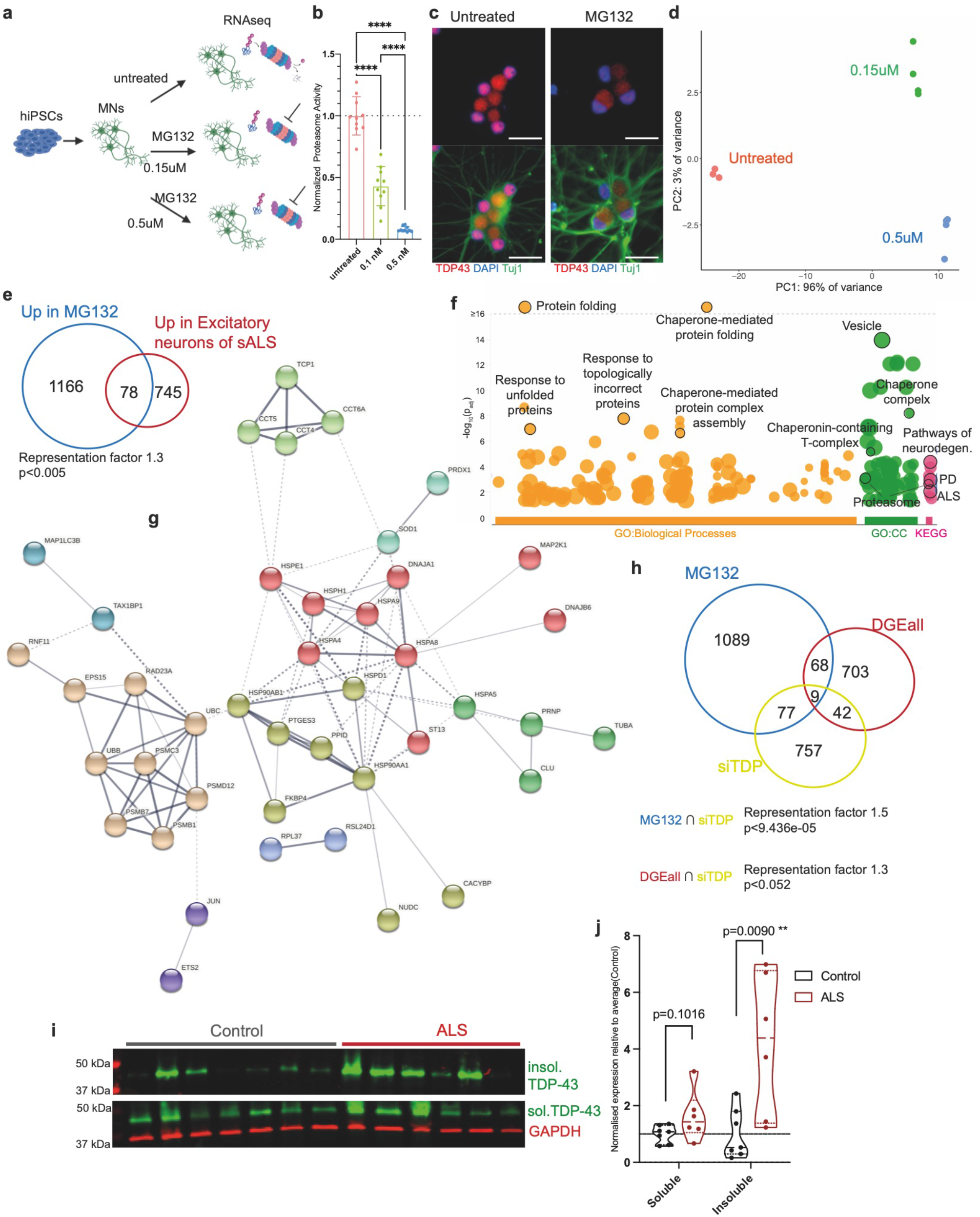
Proteostatic stress in hPSC-derived neurons resembles changes in excitatory neurons from brain of ALS patients. **a.** Diagram of neuronal differentiation from Pluripotent Stem Cells and treatment with proteasome inhibitors for bulk RNA-sequencing. **b.** Quantification of proteasome inhibition. **c.** Immunofluorescence of TDP-43 localisation after treatment. **d.** Principle Component Analysis plot showing strong effect of treatments compared to untreated controls. **e.** Venn Diagram depicting shared upregulated genes between treated hPSC-derived neurons and excitatory neurons from ALS patients. **f.** Gene Ontology analysis for shared genes in (e), highlighted terms involved in protein folding and neurodegenerative diseases (CC=Cellular Components). **g.** Protein-protein interaction network of shared genes from (d). **h.** Venn Diagram depicting shared upregulated genes in treated hPSC-derived neurons (MG132), excitatory neurons from ALS patients (DGEall) and genes misregulated in human neurons after TDP-43 siRNA from Klim et al. **i-j.** Western Blot quantification of soluble and insoluble TDP-43 from MotorCortex of ALS patients and age-matched controls.

**Extended Data Fig. 7.**
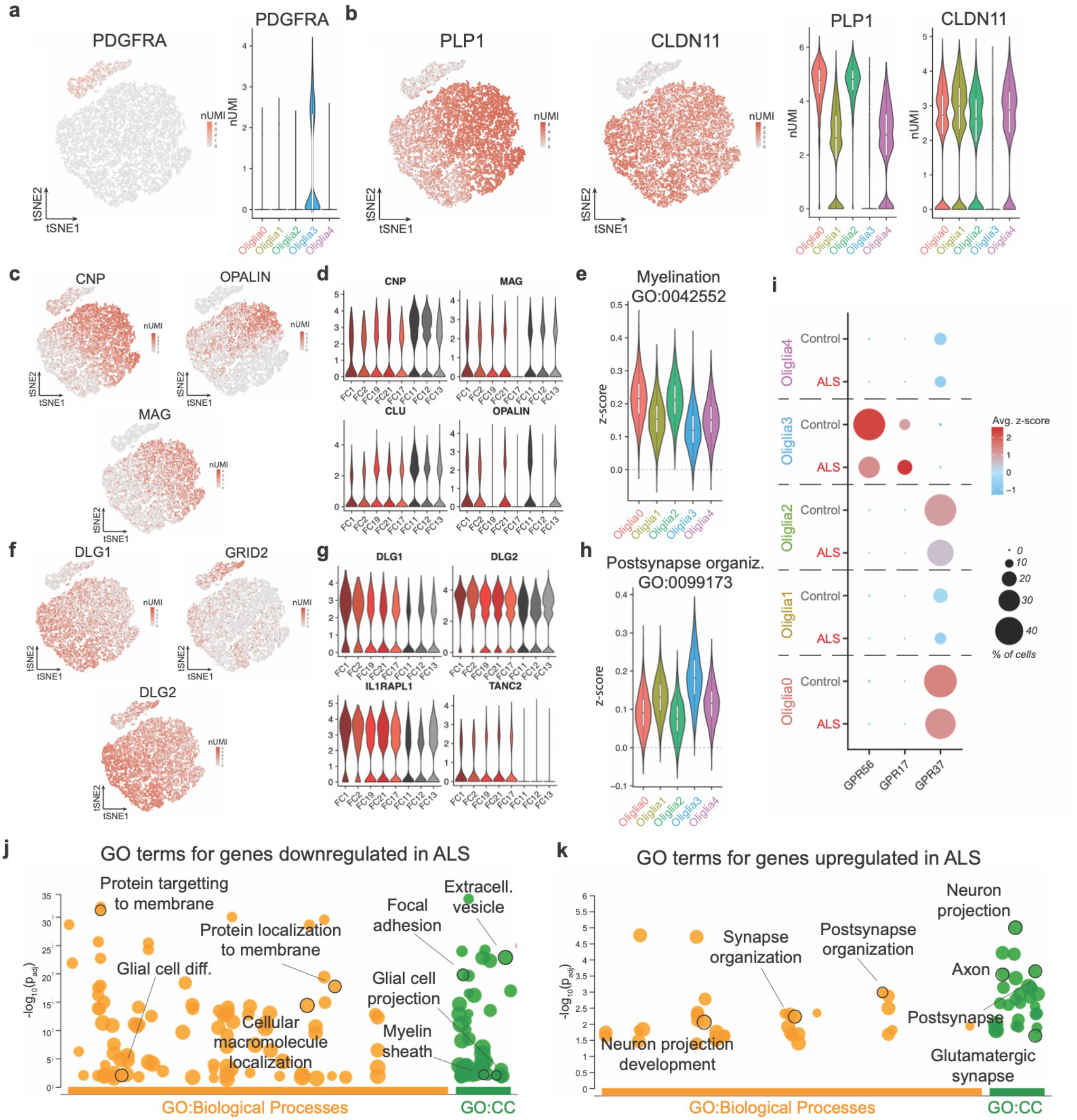
Oligodendrocytes polarize between myelinating and neuro-engaged states. **a.** *t*-SNE projection and Violin plot of markers of Oligodendrocyte Progenitor Cells (OPCs). **b.** *t*-SNE projection and Violin plots of markers of mature oligodendrocytes. **c-d.** *t*-SNE projection and Violin plots of markers of actively myelinating oligodendrocytes. **e.** Violin plots representing z-score for selected GO terms by cluster. **f-g.** *t*-SNE projection and Violin plots of markers of neuro-engaged oligodendrocytes. **h.** Violin plots representing z-score for selected GO terms by cluster. **i.** Dotplot representing genes characteristic of maturation and development of OPCs in myelinating oligodendrocytes in each subcluster split by diagnosis. **j.** GO analysis for genes downregulated in ALS oligodendrocytes, highlighted terms involved in myelination (CC=Cellular Component). **k.** GO analysis for genes upregulated in ALS oligodendrocytes, highlighted terms involved in neuro-supportive functions (CC=Cellular Component).

**Extended Data Fig. 8.**
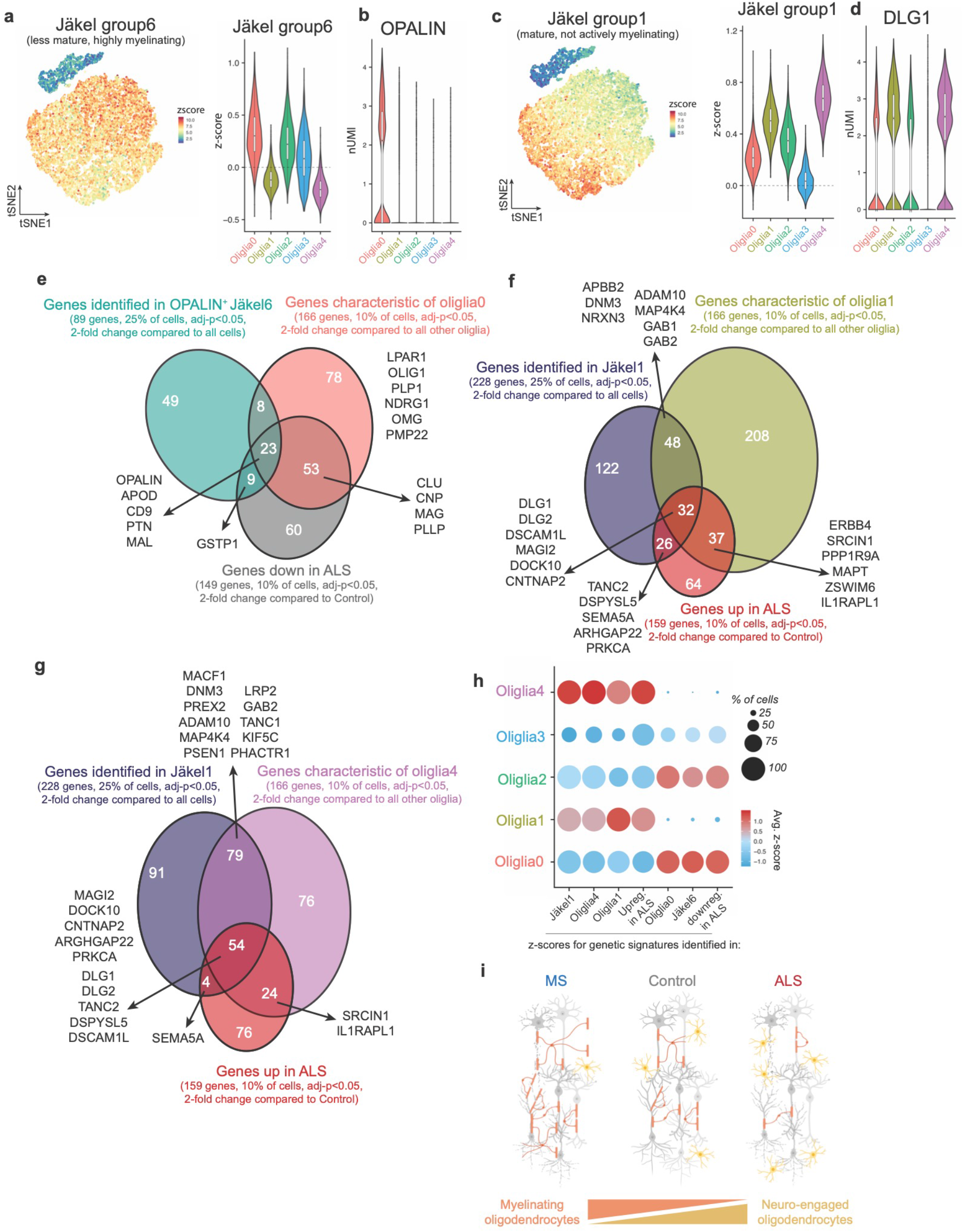
Comparison of ALS-driven changes with other studies with similar signatures disrupted in the disease. **a.** *t*-SNE projection and violin plot representing z-score for genes characteristic of highly myelinating, OPALIN^+^ oligodendrocytes in Jäkel et al. **b.** Violin plot showing OPALIN expression in our dataset. **c.** *t*-SNE projection and violin plot representing z-score for genes of mature, not-actively myelinating oligodendrocytes in Jäkel et al. **d.** Violin plot showing DLG1 expression in our dataset. **e.** Comparison of genes downregulated in oligodendroglia from ALS patients with genes characteristic of highly myelinating, OPALIN^+^ subtypes identified by this study (oliglia0) and by Jäkel et al (Jäkel6), highlighted genes are shared with GO terms shown in figures. **f,g.** Comparison of genes upregulated in oligodendroglia from ALS patients with genes characteristic of mature, lowly myelinating groups in this study (oliglia1 and 4) and by Jäkel et al (Jäkel1), highlighted genes are shared with GO terms shown in figures. **h**. Dotplot representing z-scores for the genetic signatures identified in the actively myelinating cells, the mature lowly myelinating cells and DEGs identified in this study. **i.** Diagram illustrates proposed shift of oligodendrocytes states.

**Extended Data Fig. 9.**
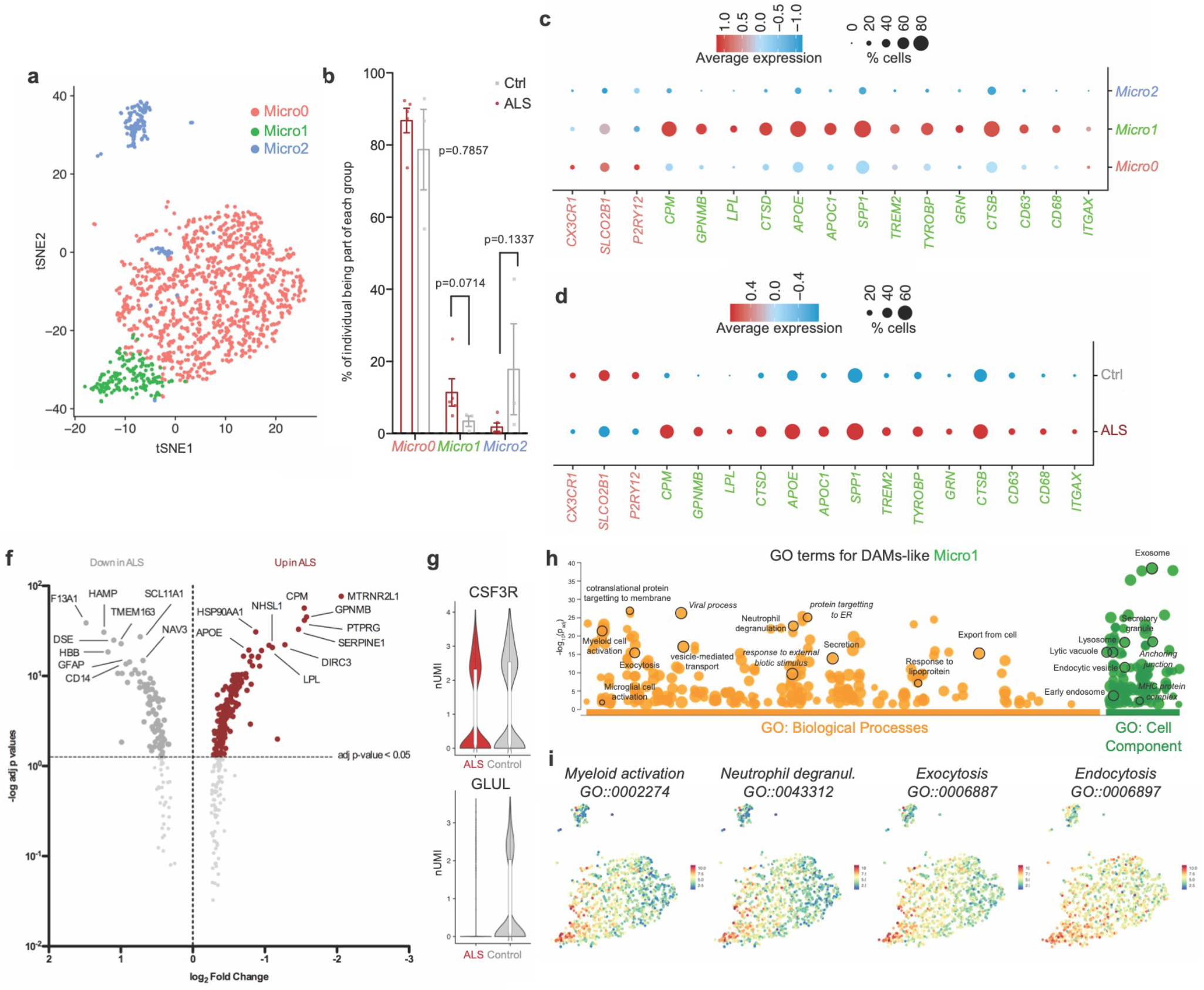
Shared features between ALS-driven changes and reactive subcluster of microglia. **a.** *t*-SNE projection of subclusters identified within microglia (Micro0 = Homeo = homeostatic, Micro1 = DAMs = Disease-associated microglia, Micro_2_ = Cycling cells)). **b.** Distribution of microglia within clusters by diagnosis. **c.** Distribution of microglia within subclusters by individual. **d.** Dotplot representing genes identified as characteristic of Homeostatic microglia and DAMs by subcluster. **e.** Dotplot representing genes identified as characteristic of Homeostatic microglia and DAMs by diagnosis. **f.** Volcano plot of statistically significant differentially expressed genes between Control and ALS microglia (top ten upregulated and top ten downregulated genes highlighted). **g**. Violin plots of representative DEGs downregulated in ALS patients of genes associated with homeostatic microglia. **h.** Gene Ontology analysis of terms associated with genes characteristic of DAMs microglia, highlighted terms playing important role in microglial biology and/or pathogenesis of the disease. **i.** *t*-SNE projections representing z-score for selected, statistically significant GO terms.

**Extended Data Fig. 10.**
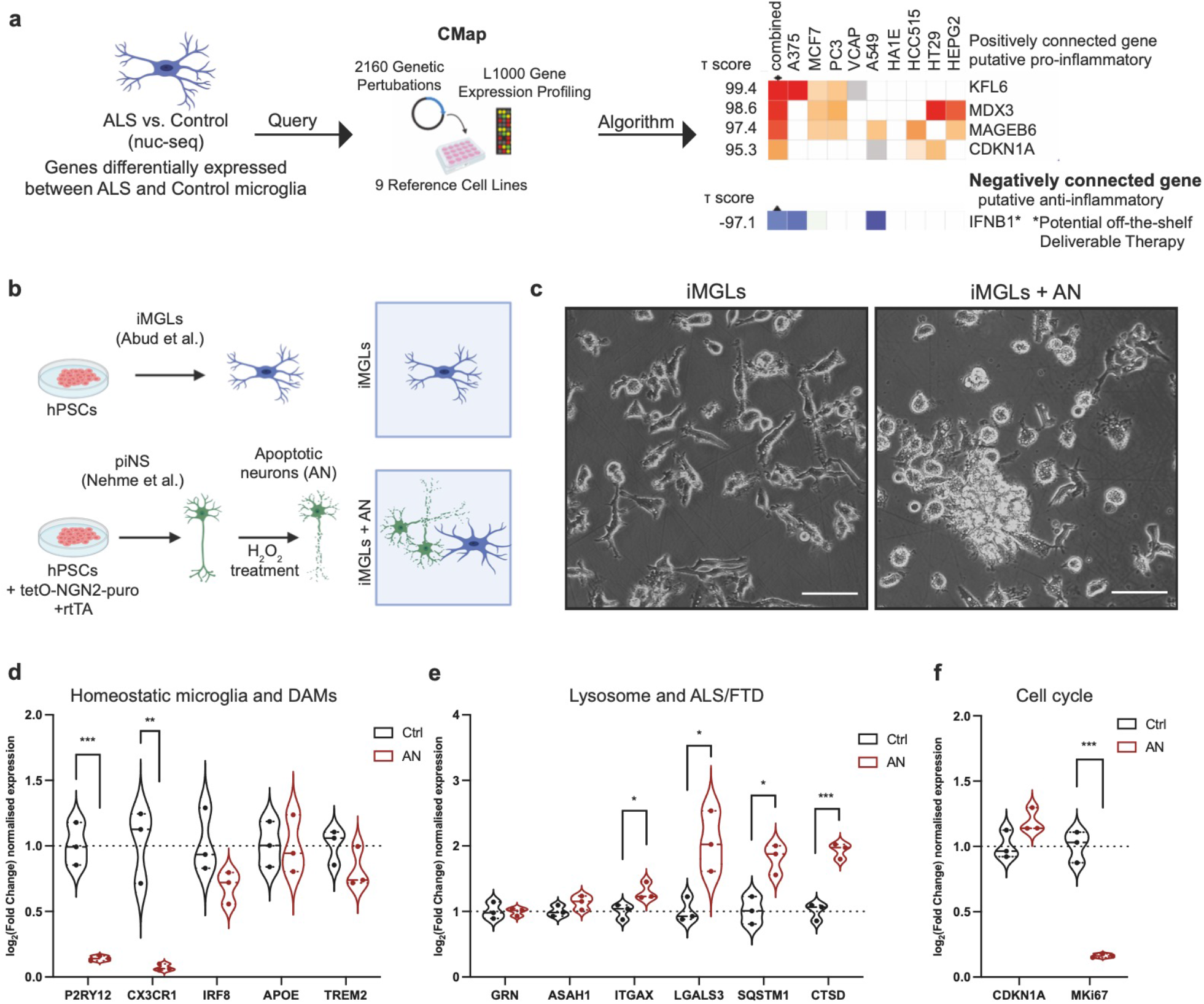
Apoptotic neurons upregulate lysosomal genes in microglia. **a.** Schematic of workflow and results from the Connectivity Map project for the genes upregulated in ALS microglia. Heatmap shows what cellular signature is most closely related to the query. **b.** Diagram of microglia and neuronal differentiation from Pluripotent Stem Cells, induction of apoptosis neurons and feeding to iMGLs. **c.** Brightfield images of untreated day 40 iMGLs and day 40 iMGLs fed apoptotic neurons for 24 hours. **d.** RT-qPCR quantification of selectedALS-FTD-associated and lysosomal genes 24h after feeding iMGLs with apoptotic neurons. **e.** RT-qPCR quantification of homeostatic and DAMs genes after feeding. **e.** RT-qPCR quantification of cell cycle-associated genes after feeding.

## Notes

### Competing Interest Statement

The authors have declared no competing interest.

### Summary of Updates

New figures and text

## References

1 Taylor, J. P., Brown, R. H., Jr. & Cleveland, D. W. Decoding ALS: from genes to mechanism. Nature 539, 197–206, doi:10.1038/nature20413 (2016).

2 Brown, R. H. & Al-Chalabi, A. Amyotrophic Lateral Sclerosis. N Engl J Med 377, 162–172, doi:10.1056/NEJMra1603471 (2017).

3 Wainger, B. J. & Lagier-Tourenne, C. Taking on the Elephant in the Tissue Culture Room: iPSC Modeling for Sporadic ALS. Cell Stem Cell 23, 466–467, doi:10.1016/j.stem.2018.09.015 (2018).

4 Prudencio, M. et al. Distinct brain transcriptome profiles in C9orf72-associated and sporadic ALS. Nat Neurosci 18, 1175–1182, doi:10.1038/nn.4065 (2015).

5 Mordes, D. A. et al. Dipeptide repeat proteins activate a heat shock response found in C9ORF72-ALS/FTLD patients. Acta Neuropathol Commun 6, 55, doi:10.1186/s40478-018-0555-8 (2018).

6 D’Erchia, A. M. et al. Massive transcriptome sequencing of human spinal cord tissues provides new insights into motor neuron degeneration in ALS. Sci Rep 7, 10046, doi:10.1038/s41598-017-10488-7 (2017).

7 Tam, O. H. et al. Postmortem Cortex Samples Identify Distinct Molecular Subtypes of ALS: Retrotransposon Activation, Oxidative Stress, and Activated Glia. Cell Rep 29, 1164–1177 e1165, doi:10.1016/j.celrep.2019.09.066 (2019).

8 Eshima, J. et al. Molecular subtypes of ALS are associated with differences in patient prognosis. Nat Commun 14, 95, doi:10.1038/s41467-022-35494-w (2023).

9 Neumann, M. et al. Ubiquitinated TDP-43 in frontotemporal lobar degeneration and amyotrophic lateral sclerosis. Science 314, 130–133, doi:10.1126/science.1134108 (2006).

10 Hammer, R. P., Jr., Tomiyasu, U. & Scheibel, A. B. Degeneration of the human Betz cell due to amyotro phic lateral sclerosis. Exp Neurol 63, 336–346, doi:10.1016/0014-4886(79)90129-8 (1979).

11 Giordana, M. T. et al. TDP-43 redistribution is an early event in sporadic amyotrophic lateral sclerosis. Brain Pathol 20, 351–360, doi:10.1111/j.1750-3639.2009.00284.x (2010).

12 Seeley, W. W. et al. Early frontotemporal dementia targets neurons unique to apes and humans. Ann Neurol 60, 660–667, doi:10.1002/ana.21055 (2006).

13 Nana, A. L. et al. Neurons selectively targeted in frontotemporal dementia reveal early stage TDP-43 pathobiology. Acta Neuropathol 137, 27–46, doi:10.1007/s00401-018-1942-8 (2019).

14 Suzuki, N. et al. The mouse C9ORF72 ortholog is enriched in neurons known to degenerate in ALS and FTD. Nat Neurosci 16, 1725–1727, doi:10.1038/nn.3566 (2013).

15 Ransohoff, R. M. How neuroinflammation contributes to neurodegeneration. Science 353, 777–783, doi:10.1126/science.aag2590 (2016).

16 Kang, S. H. et al. Degeneration and impaired regeneration of gray matter oligodendrocytes in amyotrophic lateral sclerosis. Nat Neurosci 16, 571–579, doi:10.1038/nn.3357 (2013).

17 Boillee, S. et al. Onset and progression in inherited ALS determined by motor neurons and microglia. Science 312, 1389–1392, doi:10.1126/science.1123511 (2006).

18 Schirmer, L. et al. Neuronal vulnerability and multilineage diversity in multiple sclerosis. Nature 573, 75–82, doi:10.1038/s41586-019-1404-z (2019).

19 Jakel, S. et al. Altered human oligodendrocyte heterogeneity in multiple sclerosis. Nature 566, 543–547, doi: 10.1038/s41586-019-0903-2 (2019).

20 Mathys, H. et al. Single-cell transcriptomic analysis of Alzheimer’s disease. Nature 570, 332–337, doi:10.1038/s41586-019-1195-2 (2019).

21 Zhou, Y. et al. Human and mouse single-nucleus transcriptomics reveal TREM2-dependent and TREM2-independent cellular responses in Alzheimer’s disease. Nat Med 26, 131–142, doi:10.1038/s41591-019-0695-9 (2020).

22 Leng, K. et al. Molecular characterization of selectively vulnerable neurons in Alzheimer’s disease. Nat Neurosci, doi:10.1038/s41593-020-00764-7 (2021).

23 Kamath, T. et al. Single-cell genomic profiling of human dopamine neurons identifies a population that selectively degenerates in Parkinson’s disease. Nat Neurosci 25, 588–595, doi:10.1038/s41593-022-01061-1 (2022).

24 Sadick, J. S. et al. Astrocytes and oligodendrocytes undergo subtype-specific transcriptional changes in Alzheimer’s disease. Neuron 110, 1788–1805 e1710, doi:10.1016/j.neuron.2022.03.008 (2022).

25 Grubman, A. et al. A single-cell atlas of entorhinal cortex from individuals with Alzheimer’s disease reveals cell-type-specific gene expression regulation. Nat Neurosci 22, 2087–2097, doi:10.1038/s41593-019-0539-4 (2019).

26 Lau, S. F., Cao, H., Fu, A. K. Y. & Ip, N. Y. Single-nucleus transcriptome analysis reveals dysregulation of angiogenic endothelial cells and neuroprotective glia in Alzheimer’s disease. Proc Natl Acad Sci U S A 117, 25800–25809, doi:10.1073/pnas.2008762117 (2020).

27 Gerrits, E. et al. Neurovascular dysfunction in GRN-associated frontotemporal dementia identified by single-nucleus RNA sequencing of human cerebral cortex. Nat Neurosci 25, 1034–1048, doi:10.1038/s41593-022-01124-3 (2022).

28 Macosko, E. Z. et al. Highly Parallel Genome-wide Expression Profiling of Individual Cells Using Nanoliter Droplets. Cell 161, 1202–1214, doi:10.1016/j.cell.2015.05.002 (2015).

29 Stuart, T. et al. Comprehensive Integration of Single-Cell Data. Cell 177, 1888–1902 e1821, doi:10.1016/j.cell.2019.05.031 (2019).

30 Kelley, K. W., Nakao-Inoue, H., Molofsky, A. V. & Oldham, M. C. Variation among intact tissue samples reveals the core transcriptional features of human CNS cell classes. Nat Neurosci 21, 1171–1184, doi:10.1038/s41593-018-0216-z (2018).

31 Lake, B. B. et al. Integrative single-cell analysis of transcriptional and epigenetic states in the human adult brain. Nat Biotechnol 36, 70–80, doi:10.1038/nbt.4038 (2018).

32 Tirosh, I. et al. Dissecting the multicellular ecosystem of metastatic melanoma by single-cell RNA-seq. Science 352, 189–196, doi:10.1126/science.aad0501 (2016).

33 Kunkle, B. W. et al. Genetic meta-analysis of diagnosed Alzheimer’s disease identifies new risk loci and implicates Abeta, tau, immunity and lipid processing. Nat Genet 51, 414–430, doi:10.1038/s41588-019-0358-2 (2019).

34 Jansen, I. E. et al. Genome-wide meta-analysis identifies new loci and functional pathways influencing Alzheimer’s disease risk. Nat Genet 51, 404–413, doi:10.1038/s41588-018-0311-9 (2019).

35 International Multiple Sclerosis Genetics, C. Multiple sclerosis genomic map implicates peripheral immune cells and microglia in susceptibility. Science 365, doi:10.1126/science.aav7188 (2019).

36 Paolicelli, R. C. et al. Microglia states and nomenclature: A field at its crossroads. Neuron 110, 3458–3483, doi:10.1016/j.neuron.2022.10.020 (2022).

37 Dolan, M. J. et al. A resource for generating and manipulating human microglial states in vitro. bioRxiv [PREPRINT], doi:doi.org/10.1101/2022.05.02.490100 (2022).

38 Arlotta, P. et al. Neuronal subtype-specific genes that control corticospinal motor neuron development in vivo. Neuron 45, 207–221, doi:10.1016/j.neuron.2004.12.036 (2005).

39 Ozdinler, P. H. et al. Corticospinal motor neurons and related subcerebral projection neurons undergo early and specific neurodegeneration in hSOD1G(9)(3)A transgenic ALS mice. JNeurosci 31, 4166–4177, doi:10.1523/JNEUROSCI.4184-10.2011 (2011).

40 Tsang, Y. M., Chiong, F., Kuznetsov, D., Kasarskis, E. & Geula, C. Motor neurons are rich in non-phosphorylated neurofilaments: cross-species comparison and alterations in ALS. Brain Res 861, 45–58, doi:10.1016/s0006-8993(00)01954-5 (2000).

41 Bakken, T. E. et al. Comparative cellular analysis of motor cortex in human, marmoset and mouse. Nature 598, 111–119, doi:10.1038/s41586-021-03465-8 (2021).

42 Guerra San Juan, I. et al. Loss of mouse Stmn2 function causes motor neuropathy. Neuron, doi:10.1016/j.neuron.2022.02.011 (2022).

43 Rivara, C. B., Sherwood, C. C., Bouras, C. & Hof, P. R. Stereologic characterization and spatial distribution patterns of Betz cells in the human primary motor cortex. Anat Rec A Discov Mol Cell Evol Biol 270, 137–151, doi:10.1002/ar.a.10015 (2003).

44 Hodge, R. D. et al. Transcriptomic evidence that von Economo neurons are regionally specialized extratelencephalic-projecting excitatory neurons. Nat Commun 11, 1172, doi:10.1038/s41467-020-14952-3 (2020).

45 Cobos, I. & Seeley, W. W. Human von Economo neurons express transcription factors associated with Layer V subcerebral projection neurons. Cereb Cortex 25, 213–220, doi:10.1093/cercor/bht219 (2015).

46 Zeng, H. et al. Large-scale cellular-resolution gene profiling in human neocortex reveals species-specific molecular signatures. Cell 149, 483–496, doi:10.1016/j.cell.2012.02.052 (2012).

47 Maynard, K. R. et al. Transcriptome-scale spatial gene expression in the human dorsolateral prefrontal cortex. Nat Neurosci 24, 425–436, doi:10.1038/s41593-020-00787-0 (2021).

48 Velmeshev, D. et al. Single-cell genomics identifies cell type-specific molecular changes in autism. Science 364, 685–689, doi:10.1126/science.aav8130 (2019).

49 Braak, H., Ludolph, A. C., Neumann, M., Ravits, J. & Del Tredici, K. Pathological TDP-43 changes in Betz cells differ from those in bulbar and spinal alpha-motoneurons in sporadic amyotrophic lateral sclerosis. Acta Neuropathol 133, 79–90, doi:10.1007/s00401-016-1633-2 (2017).

50 Porta, S. et al. Patient-derived frontotemporal lobar degeneration brain extracts induce formation and spreading of TDP-43 pathology in vivo. Nat Commun 9, 4220, doi:10.1038/s41467-018-06548-9 (2018).

51 van Rheenen, W. et al. Common and rare variant association analyses in amyotrophic lateral sclerosis identify 15 risk loci with distinct genetic architectures and neuron-specific biology. Nat Genet 53, 1636–1648, doi:10.1038/s41588-021-00973-1 (2021).

52 Saez-Atienzar, S. et al. Genetic analysis of amyotrophic lateral sclerosis identifies contributing pathways and cell types. Sci Adv 7, doi:10.1126/sciadv.abd9036 (2021).

53 Baxi, E. G. et al. Answer ALS, a large-scale resource for sporadic and familial ALS combining clinical and multi-omics data from induced pluripotent cell lines. Nat Neurosci 25, 226–237, doi:10.1038/s41593-021-01006-0 (2022).

54 Giacomelli, E. et al. Human stem cell models of neurodegeneration: From basic science of amyotrophic lateral sclerosis to clinical translation. Cell Stem Cell 29, 11–35, doi:10.1016/j.stem.2021.12.008 (2022).

55 Limone, F., Klim, J. R. & Mordes, D. A. Pluripotent stem cell strategies for rebuilding the human brain. Front Aging Neurosci 14, 1017299, doi:10.3389/fnagi.2022.1017299 (2022).

56 Yu, H. et al. HSP70 chaperones RNA-free TDP-43 into anisotropic intranuclear liquid spherical shells. Science 371, doi:10.1126/science.abb4309 (2021).

57 Gwon, Y. et al. Ubiquitination of G3BP1 mediates stress granule disassembly in a context-specific manner. Science 372, eabf6548, doi:10.1126/science.abf6548 (2021).

58 Klim, J. R. et al. ALS-implicated protein TDP-43 sustains levels of STMN2, a mediator of motor neuron growth and repair. Nat Neurosci 22, 167–179, doi:10.1038/s41593-018-0300-4 (2019).

59 Limone, F. et al. Efficient generation of lower induced motor neurons by coupling Ngn2 expression with developmental cues. Cell Rep, 111896, doi:10.1016/j.celrep.2022.111896 (2022).

60 Tomassy, G. S. et al. Distinct profiles of myelin distribution along single axons of pyramidal neurons in the neocortex. Science 344, 319–324, doi:10.1126/science.1249766 (2014).

61 Giera, S. et al. The adhesion G protein-coupled receptor GPR56 is a cell-autonomous regulator of oligodendrocyte development. Nat Commun 6, 6121, doi:10.1038/ncomms7121 (2015).

62 Yang, H. J., Vainshtein, A., Maik-Rachline, G. & Peles, E. G protein-coupled receptor 37 is a negative regulator of oligodendrocyte differentiation and myelination. Nat Commun 7, 10884, doi:10.1038/ncomms10884 (2016).

63 Keren-Shaul, H. et al. A Unique Microglia Type Associated with Restricting Development of Alzheimer’s Disease. Cell 169, 1276–1290 e1217, doi:10.1016/j.cell.2017.05.018 (2017).

64 de Boer, A. S. et al. Genetic validation of a therapeutic target in a mouse model of ALS. Sci Transl Med 6, 248ra104, doi:10.1126/scitranslmed.3009351 (2014).

65 Zhang, Y. et al. The C9orf72-interacting protein Smcr8 is a negative regulator of autoimmunity and lysosomal exocytosis. Genes Dev 32, 929–943, doi:10.1101/gad.313932.118 (2018).

66 Burberry, A. et al. C9orf72 suppresses systemic and neural inflammation induced by gut bacteria. Nature 582, 89–94, doi:10.1038/s41586-020-2288-7 (2020).

67 Subramanian, A. et al. A Next Generation Connectivity Map: L1000 Platform and the First 1,000,000 Profiles. Cell 171, 1437–1452 e1417, doi:10.1016/j.cell.2017.10.049 (2017).

68 Abud, E. M. et al. iPSC-Derived Human Microglia-like Cells to Study Neurological Diseases. Neuron 94, 278–293 e279, doi:10.1016/j.neuron.2017.03.042 (2017).

69 Masuda, T. et al. Spatial and temporal heterogeneity of mouse and human microglia at single-cell resolution. Nature 566, 388–392, doi:10.1038/s41586-019-0924-x (2019).

70 Blum, J. A. & Gitler, A. D. Singling out motor neurons in the age of single-cell transcriptomics. Trends Genet, doi:10.1016/j.tig.2022.03.016 (2022).

71 Farhan, S. M. K. et al. Exome sequencing in amyotrophic lateral sclerosis implicates a novel gene, DNAJC7, encoding a heatshock protein. Nat Neurosci 22, 1966–1974, doi:10.1038/s41593-019-0530-0 (2019).

72 Melamed, Z. et al. Premature polyadenylation-mediated loss of stathmin-2 is a hallmark of TDP-43-dependent neurodegeneration. Nat Neurosci 22, 180–190, doi:10.1038/s41593-018-0293-z (2019).

73 Otero-Garcia, M. et al. Molecular signatures underlying neurofibrillary tangle susceptibility in Alzheimer’s disease. Neuron 110, 2929–2948 e2928, doi:10.1016/j.neuron.2022.06.021 (2022).

74 Pineda, S. S. et al. Single-cell profiling of the human primary motor cortex in ALS and FTLD. bioRxiv [PREPRINT], doi:doi.org/10.1101/2021.07.07.451374 (2021).

75 Yadav, A. et al. A Cellular Taxonomy of the Adult Human Spinal Cord. bioRxiv [PREPRINT], doi:doi.org/10.1101/2022.03.25.485808 (2022).

76 Falcao, A. M. et al. Disease-specific oligodendrocyte lineage cells arise in multiple sclerosis. Nat Med 24, 1837–1844, doi:10.1038/s41591-018-0236-y (2018).

77 Hughes, A. N. & Appel, B. Oligodendrocytes express synaptic proteins that modulate myelin sheath formation. Nat Commun 10, 4125, doi:10.1038/s41467-019-12059-y (2019).

78 Genc, B. et al. Apical dendrite degeneration, a novel cellular pathology for Betz cells in ALS. Sci Rep 7, 41765, doi:10.1038/srep41765 (2017).

79 Wainger, B. J. et al. Effect of Ezogabine on Cortical and Spinal Motor Neuron Excitability in Amyotrophic Lateral Sclerosis: A Randomized Clinical Trial. JAMA Neurol 78, 186–196, doi:10.1001/jamaneurol.2020.4300 (2021).

80 O’Rourke, J. G. et al. C9orf72 is required for proper macrophage and microglial function in mice. Science 351, 1324–1329, doi:10.1126/science.aaf1064 (2016).

81 McCauley, M. E. et al. C9orf72 in myeloid cells suppresses STING-induced inflammation. Nature 585, 96–101, doi:10.1038/s41586-020-2625-x (2020).

82 Ennerfelt, H. et al. SYK coordinates neuroprotective microglial responses in neurodegenerative disease. Cell 185, 4135–4152 e4122, doi:10.1016/j.cell.2022.09.030 (2022).

83 Pandya, V. A. & Patani, R. Decoding the relationship between ageing and amyotrophic lateral sclerosis: a cellular perspective. Brain 143, 1057–1072, doi:10.1093/brain/awz360 (2020).

84 Marsh, S. E. et al. Dissection of artifactual and confounding glial signatures by single-cell sequencing of mouse and human brain. Nat Neurosci 25, 306–316, doi:10.1038/s41593-022-01022-8 (2022).

## References

85 Krienen, F. M. et al. Innovations present in the primate interneuron repertoire. Nature 586, 262–269, doi:10.1038/s41586-020-2781-z (2020).

86 Saunders, A. et al. Molecular Diversity and Specializations among the Cells of the Adult Mouse Brain. Cell 174, 1015–1030 e1016, doi:10.1016/j.cell.2018.07.028 (2018).

87 Mitchell, J. M. et al. Mapping genetic effects on cellular phenotypes with “cell villages”. bioRxiv [PREPRINT], doi:doi.org/10.1101/2020.06.29.174383 (2020).

88 Limone, F. et al. Single-nucleus sequencing reveals enriched expression of genetic risk factors sensitises Motor Neurons to degeneration in ALS. bioRxiv [PREPRINT], doi:doi.org/10.1101/2021.07.12.452054 (2021).

89 Rapino, F. et al. Small-molecule screen reveals pathways that regulate C4 secretion in stem cell-derived astrocytes. Stem Cell Reports 18, 237–253, doi:10.1016/j.stemcr.2022.11.018 (2023).

90 Finak, G. et al. MAST: a flexible statistical framework for assessing transcriptional changes and characterizing heterogeneity in single-cell RNA sequencing data. Genome Biol 16, 278, doi:10.1186/s13059-015-0844-5 (2015).

91 Hao, Y. et al. Integrated analysis of multimodal single-cell data. Cell 184, 3573–3587 e3529, doi:10.1016/j.cell.2021.04.048 (2021).

92 Raudvere, U. et al. g:Profiler: a web server for functional enrichment analysis and conversions of gene lists (2019 update). Nucleic Acids Res 47, W191–W198, doi:10.1093/nar/gkz369 (2019).

93 Koopmans, F. et al. SynGO: An Evidence-Based, Expert-Curated Knowledge Base for the Synapse. Neuron 103, 217–234 e214, doi:10.1016/j.neuron.2019.05.002 (2019).

94 Szklarczyk, D. et al. STRING v11: protein-protein association networks with increased coverage, supporting functional discovery in genome-wide experimental datasets. Nucleic Acids Res 47, D607–D613, doi:10.1093/nar/gky1131 (2019).

95 Subramanian, A. et al. Gene set enrichment analysis: a knowledge-based approach for interpreting genome-wide expression profiles. Proc Natl Acad Sci U S A 102, 15545–15550, doi:10.1073/pnas.0506580102 (2005).

96 Nehme, R. et al. Combining NGN2 Programming with Developmental Patterning Generates Human Excitatory Neurons with NMDAR-Mediated Synaptic Transmission. Cell Rep 23, 2509–2523, doi:10.1016/j.celrep.2018.04.066 (2018).

97 Pietilainen, O. et al. Astrocytic cell adhesion genes linked to schizophrenia correlate with synaptic programs in neurons. Cell Rep 42, 111988, doi:10.1016/j.celrep.2022.111988 (2023).

98 Fukuda, A. et al. De novo DNA methyltransferases DNMT3A and DNMT3B are essential for XIST silencing for erosion of dosage compensation in pluripotent stem cells. Stem Cell Reports 16, 2138–2148, doi:10.1016/j.stemcr.2021.07.015 (2021).

99 Hao, J. et al. Loss of TBK1 activity leads to TDP-43 proteinopathy through lysosomal dysfunction in human motor neurons. bioRxiv [PREPRINT], doi:doi.org/10.1101/2021.10.11.464011 (2021).

